# RNA methylation controls stress granule formation and function during erythropoiesis

**DOI:** 10.1101/2024.11.07.622459

**Authors:** Rajesh Gunage, Shuibin Lin, David Wiley, Avik Choudhuri, Mackenzie Smith, Tianxiao Han, Arish Shah, Steven Coyne, Katherine Koczirka, Kenny Zhi Ming Chen, Chunjie Yu, Kyle Drake, Xin Yang, Song Yang, Yi Zhou, Daniel Bauer, Zhijian Qian, Eliezer Calo, Richard I. Gregory, Leonard I. Zon

## Abstract

Stress granules (SGs) are crucial in RNA regulation, affecting cell fate and function. SGs contain RNAs, some of which can be methylated. We studied m^6^A RNA modifications during the human CD34^+^ HSPCs (hCD34^+^) differentiating into erythroid cells and found that mRNAs encoding many erythroid-specific proteins had decreased methylation during differentiation. Increased levels ALKBH5 demethylase during erythropoiesis controls the levels of the 3’UTR methylation of these mRNAs. hCD34^+^ carrying ALKBH5 mutations demonstrated a block in erythropoiesis, and mass-spectrometry studies of the mutant cells showed decreased levels of SG proteins, including the core granule protein ATXN2. ALKBH5 directly regulates the methylation of the mRNA of *ATXN2*. *ATXN2* overexpression accelerated the erythroid differentiation of HSPCs and rescued the erythroid differentiation of *ALKBH5* mutant cells. Very few SGs are found in normal human erythroid progenitors. SGs accumulated substantially in *ALKBH5* mutant cells, and surprisingly overexpression of *ATNX2* reduced SG numbers to normal. Polysome analysis demonstrated m^6^A-modified RNAs to be enriched in the pre-polysome fractions that were less translated. This work establishes a mechanism by which during stress, ATXN2 facilitates the release of SG-stored m^6^A-modified RNAs including erythroid-specific and SG-enriched RNAs that are loaded onto functional ribosomes, allowing better translation and accelerated erythroid differentiation during stress.

## Introduction

m^6^A RNA modification has been shown to be a major regulator of cell fate decisions (Ma et al., 2018; Zhang et al., 2017). m^6^A modifications are co-transcriptional, impacting RNA stability and translation. The modification regulates RNA binding to proteins and cytoplasmic localization (Das et al., 2022; Fu and Zhuang, 2020; Khong et al., 2022). Ribonucleoprotein (RNP) granules coordinate RNA metabolism in a hub of proteins and RNA. Several proteins form a common subset of RNP granules (Anderson and Kedersha, 2009; Ripin and Parker, 2022). Stress granules, or cytoplasmic RNP assemblies, form a hub for RNA metabolism controlling RNA fate via RNA storage, stability, and translation (Bakthavachalu et al., 2018; Wolozin and Ivanov, 2019; Yang et al., 2020). Stress granules are membrane-less organelles composed of RNA-binding proteins such (RBPs) as ATXN2, G3BP1/2, PABP, Tia-1, and TDP-43, and stored RNAs (Inagaki et al., 2020; Protter and Parker, 2016). Dysregulation of SG RBPs affects cell function and health. Mutations in SG core components lead to diseases via altered ‘ribostasis.’ (Ramaswami et al., 2013; Sproviero et al., 2017; Vanderweyde et al., 2012).

Stress granules (SGs) are critical in RNA regulation during the stress response. SGs are reported to occur in numerous cell types and are directly involved in the transcriptome regulation (Bakthavachalu et al., 2018; Tauber et al., 2020). SG core proteins such as ATAXIN2 (ATXN-2) gene mutations lead to diseases such as Ataxia and other neuromuscular disorders. Disease mutations lead to altered SGs composition, size and number due to changes in the protein stability and properties (Damrath et al., 2012; Kahle et al., 2013; Sproviero et al., 2017). SGs assemble under stress conditions, but how the individual core proteins are regulated is unknown at RNA and protein levels (Vanderweyde et al., 2012; Yang et al., 2020).

RNA regulation is vital for stem cell function and lineage commitment (Geula et al., 2015; Yang et al., 2020). m^6^A RNA modifications are the most abundant and dynamic RNA modifications crucial in cell fate and function. m^6^A modifications are added co-transcriptionally by PCIF1, Mettl3/Mettl14, and WTAP (Akichika et al., 2019; Cowling, 2019; Geula et al., 2015; Roundtree et al., 2017; Vu et al., 2017). FTO and Alkbh5 RNA demethylases remove the m^6^A modification (Mauer et al., 2017; Zaccara et al., 2019; Zheng et al., 2013). m^6^A-mediated gene regulation plays a key role in blood development and the leukemia (Gao et al., 2021; Shen et al., 2018, 2015). ALKBH5 RNA demethylase-mediated m^6^A regulation is key to normal spermatogenesis, and ALKBH5 loss of function inhibits leukemogenesis due to deregulation of TACC3, AXL oncogenes (Cheng et al., 2020; Shen et al., 2020; Tang et al., 2017; Wang et al., 2020).

Here, we used human bone marrow immobilized CD34^+^ (hCD34^+^) hematopoietic stem and progenitor stem cells to study SG core component regulation via the m^6^A modification. By studying human erythropoiesis in vitro, we show numerous transcripts to be m^6^A modified, including erythroid-specific genes and the SG core component gene transcripts. We demonstrate 3’UTRs of *ATXN2*, *G3BP1/2*, *Tia-1*, and other SGs RNA binding proteins (RBP) mRNAs are modified with m^6^A. ALKBH5 demethylase regulates m^6^A levels on these transcripts and directly affects the protein production of core SG protein ATXN2. Gene knockdown or knockout of *ATXN2* results in SGs aggregation containing m^6^A RNA and a block in erythropoiesis. Restoring ATXN2 levels alleviates massive cytoplasmic RNA stress granules and enhances the translation of m^6^A-marked RNAs. Overexpression of *ATXN2* promoted erythropoiesis in hCD34+ cells, revealing its function in stress-induced blood formation and regeneration.

## Results

### RNA m^6^A methylation decreases during human erythropoiesis

To study m^6^A modifications in mRNAs during hematopoietic differentiation, hCD34^+^ stem and progenitor cells were differentiated into erythroid cells (Choudhuri et al., 2020; Giarratana et al., 2011) using SCF, IL-3, and Epo and transcriptome-wide temporal m^6^A profiling was performed (Fig. 1a-e and supplementary Fig. 1a). We analyzed levels of m^6^A-marked transcripts using MeRIP sequencing (methylated RNA immunoprecipitation sequencing) (Dominissini et al., 2013). hCD34+ cell-derived erythroid cells at proerythroblast stages have ~4800 of transcripts with m^6^A modifications. Red blood cell fate-specific genes such as *HBB* and *TFRC* were preferentially m^6^A modified (Fig. 1 a-e). MeRIP sequencing and analysis of later differentiation stages showed reduced m^6^A levels at a transcriptome-wide level. m^6^A on erythropoiesis-specific gene transcripts, such as *HBB* and *TFRC*, showed a significant reduction in m^6^A-methylation at the 3’ end as the differentiation progressed (Fig 1a, b and d) (*p-Values< 0.01, ** p-Value<0.001) (Table 1). The house-keeping gene *GAPDH* mRNA m^6^A level showed no such trend (Fig. 1c, d). The m^6^A modification was distributed in 5’UTR (11%), exon (46%), intron (15%), and 3’ UTR (28%), respectively (supplementary Fig. 1b). A transcriptome-wide motif analysis of m^6^A locations identified U/AGGAC as the consensus motif, consistent with a motif reported previously (Fig. 1e) (Dominissini et al., 2013). The m^6^A levels of transcriptome on days 7 (proerythroblasts) and 10 (late erythroid-polychromatic erythroblast) of erythropoiesis showed progressive demethylation (Fig. 1f, g). Transcriptome-wide analysis of the m^6^A peaks showed 4368 unique peaks to be hypomethylated from the proerythroblasts stage to the late erythroid stage (Fig. 1g). We performed pathway analysis of these peaks, which showed enrichment of the genes involved in the RNA metabolism and RNA-protein complex biogenesis (Fig. 1h). Similar analysis of hyper-methylated unique 576 peaks corresponded to leukocyte development and T-cell processes. (Fig. 1g and supplementary Fig. 1c). We confirmed these findings using K562 erythroleukemia cells differentiated under hemin induction (Addya et al., 2004). mRNAs of key erythroid genes, *Tal1* and *TFRC*, were marked with m^6^A, and their 3’ UTR m^6^A level showed a reduction during erythroid differentiation (Supplementary Fig. 1d, e, and g). On the other hand, *RPL11*, a ribosomal gene, was also m^6^A labeled but showed no increased demethylation in the process (Supplementary Fig. 1f and g.). K562 cells showed a clear demethylation trend as observed in the case of hCD34+ cells and from D1 to D3 post hemin induction, a majority of differential m^6^A peak (1185 peaks out of 1522 peaks) were hypo methylated (Supplementary Fig. 1h). In summary, MeRIP-seq analysis of hCD34+ generated erythroid cells and K562 differentiated erythroid cells showed downregulation of m^6^A levels on the mRNAs of the key erythropoiesis and RNA metabolism genes.

**Fig. 1.**
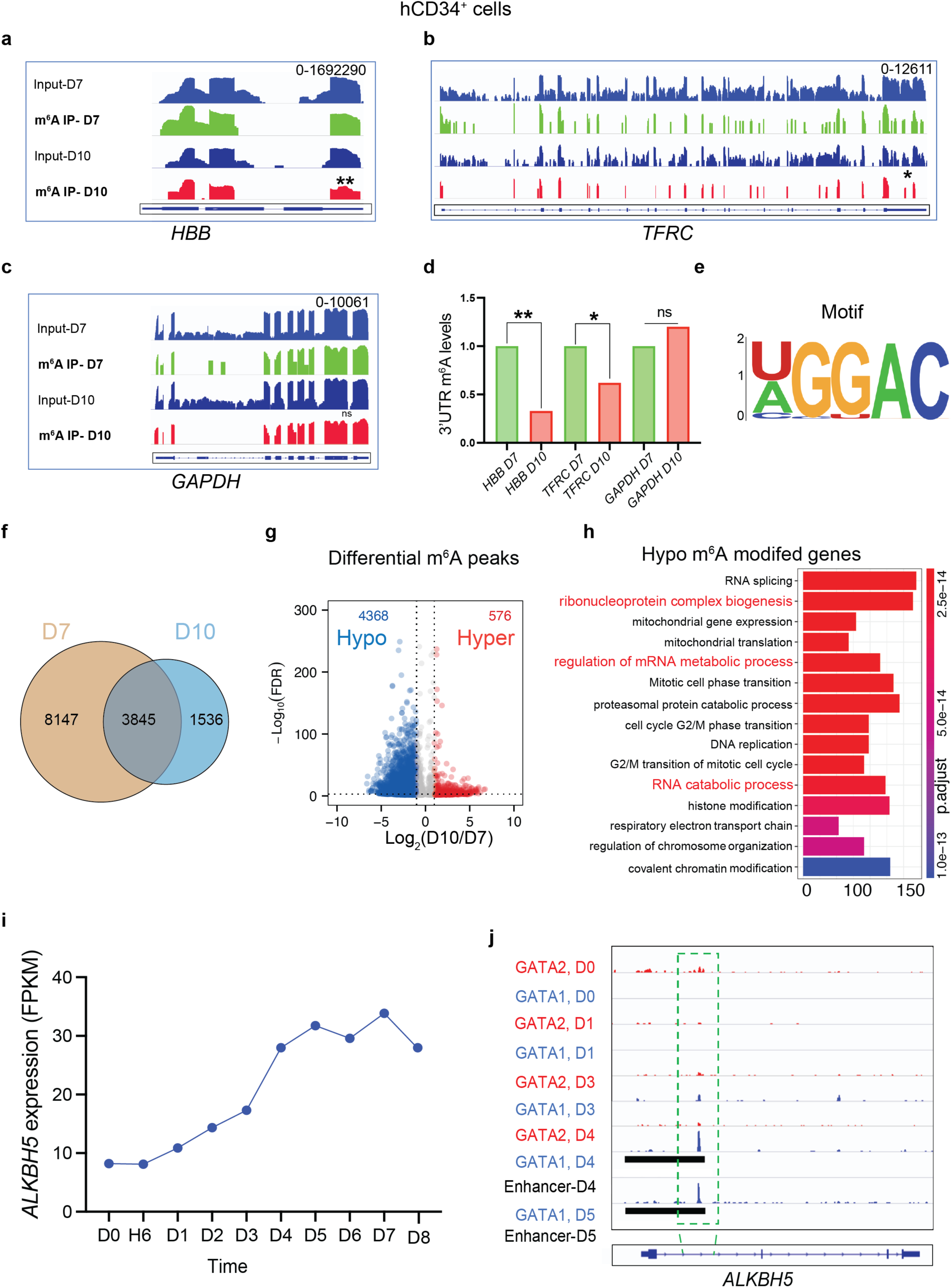
RNA m^6^A methylation decreases during human erythropoiesis. **a-h**, m^6^A analysis of human CD34+ cells during erythropoiesis. hCD34+ cells were subjected to invitro erythropoiesis, and poly-A tail RNAs were subjected to m^6^A pull down using anti-m^6^A antibody and sequencing. Blue plots represent the total RNA input sequence. Green and red plots represent m^6^A methylation marks along the gene for (Proerythroblasts) day 7 and (late erythroid-polychromatic erythroblast) day 10, respectively. Erythropoiesis-specific genes (**a**) *HBB*, (**b**) *TFRC*, and (**c**) *GAPDH* shows m^6^A modifications along gene body and 3’UTR. **d**, Quantification of m^6^A levels at 3’UTR of genes *HBB*, *TFRC*, and *GAPDH.* **e**, Transcriptome-wide analysis identified consensus motif U/AGGAC as an m^6^A modification site. **f**, Venn diagram showing the overlap of m^6^A peaks identified in mRNA of CD34+ cells at Day 7 and Day 10. **g**, Differential m^6^A peaks during erythropoiesis. Blue dots represent decreased m^6^A peaks (hypo-m^6^A), and red dots represent increased m^6^A peaks (hyper-m^6^A). Grey dots correspond to m^6^A peaks with no significant change. **h**, GO terms of genes with hypo m^6^A peaks. **i**, *ALKBH5* transcript levels during hCD34+ cell erythropoiesis. **j**, Enhancer analysis showing GATA1 and GATA2 binding at *ALKBH5* locus during hCD34+ cell erythropoiesis. p-values ** <0.001, ** <0.01 from unpaired t-test. H-hour, D-Day.

**Table 1.** hCD34^+^ cells m^6^A analysis for *HBB*, *TFRC*, and *GAPDH*.

| Gene Symbol |  | Input-5'end | m6A Ip-5' end | Input-3'end | m6A Ip-3' end | Ratio of (3' m6A)/(5' m6A) | Normalised values |
| --- | --- | --- | --- | --- | --- | --- | --- |
| hCD34+ cell |  |  |  |  |  |  |  |
| HBB D7 |  | 222490 | 1692450 | 332049 | 39546 | 63.87 | 1.00 |
| HBB D10 |  | 13357 | 18340 | 12311 | 799 | 21.16 | 0.33 |
| TFRC D7 |  | 17 | 484 | 1246 | 223 | 8.07 | 1.00 |
| TFRC D10 |  | 16 | 205 | 4396 | 543 | 5.26 | 0.65 |
| GAPDH D7 |  | 2992 | 758 | 7239 | 4922 | 0.41 | 1.00 |
| GAPDH D10 |  | 2811 | 305 | 4470 | 1610 | 0.33 | 0.81 |
| K562 Cell |  |  |  |  |  |  |  |
| TFRC D1 |  | 128 | 926 | 160 | 9 | 1.53 | 1.00 |
| TFRC D3 |  | 37 | 442 | 755 | 189 | 0.57 | 0.37 |
| Tal1 D1 |  | 84 | 258 | 31 | 780 | 0.12 | 1.00 |
| Tal1 D3 |  | 29 | 29 | 5 | 446 | 0.01 | 0.09 |
| RPL11 D1 |  | 14292 | 7052 | 9551 | 23500 | 0.70 | 1.00 |
| RPL11 D3 |  | 8332 | 23285 | 6712 | 103000 | 0.64 | 0.91 |

To define the enzymes that confer or remove m^6^A modification on 3’ UTRs, we examined the expression of *FTO* and *ALKBH5* RNA demethylases during erythroid differentiation. *ALKBH5* mRNA levels increased continuously more than threefold during hCD34^+^ cell erythropoiesis from D0 to D8 (Fig. 1i). *FTO* was not changed in expression during erythropoiesis (Supplementary Fig. 1i). We previously had undertaken ChIP-seq analysis of two master transcriptional regulators of erythropoiesis, GATA-1 and GATA-2 during hCD34+ differentiation (Choudhuri et al., 2020). GATA1-binding revealed enhancer enrichment at the *ALKBH5* RNA demethylase locus and an increased GATA1 binding at the *ALKBH5* locus from D0 to D6 of erythropoiesis (Fig. 1j), consistent with erythroid-specific enhancers in the *ALKBH5* locus. ALKBH5 catalyzes the demethylation of m^6^A marks on mRNAs and is a key component of transcriptome-wide RNA regulation via m^6^A marks (Ensfelder et al., 2018; Zheng and He, 2015). ALKBH5 demethylates m^6^A on mRNAs post-transcriptionally and regulates the cell fate (Cheng et al., 2020; Shen et al., 2020; Wang et al., 2020). We performed immunostaining of hCD34^+^ cells during erythropoiesis using an anti-ALKBH5 antibody. As shown in Supplementary Fig. 1j on day 0, ALKBH5 was primarily localized to the nucleus. For erythroid differentiation, D3-4 and later, ALKBH5 localized mainly to the cytoplasm, indicating a cytoplasmic function (Supplement Fig. 1j). In summary, ALKBH5 upregulation during erythropoiesis suggested m^6^A modification plays a role in erythropoiesis during CD34+ progenitor differentiation.

### *ALKBH5* regulates human CD34^+^ cells, zebrafish, and murine erythropoiesis

To address the role of *ALKBH5* in human erythropoiesis, we combined CRISPR Cas9 editing with the Methocult colony formation assay (Petzer et al., 1996; Tusi et al., 2018). The Methocult media includes SCF, IL-3, IL-6, EPO, G-CSF, and GM-CSF that supports optimal growth of erythroid progenitor cells (BFU-E and CFU-E), granulocyte-macrophage progenitor cells (CFU-GM, CFU-G, and CFU-M), and multipotential granulocyte, erythroid, macrophage, and megakaryocyte progenitor cells (CFU-GEMM). A Methocult assay of hCD34^+^ cells with *ALKBH5* gRNA and Cas9 mRNA showed a significant reduction in the number and size of the surviving erythroid colonies compared to the control gRNA- and Cas9-treated cells (Fig. 2a, b, t-test **p-value< 0.001). We achieved efficient *ALKBH5*-CRISPR/Cas9 editing efficiency in the hCD34+ cells, as measured by ICE Analysis (Synthego), with nearly 90% editing (Expt.1-80% deletions, 20% insertions, and Expt.2-67% deletions, 17% insertions) (Supplementary Fig. 2a). The number of CFU-E and BFU-E in the ALKBH5 deficient CD34+ cells were significantly more reduced than that of the myeloid G/M/GM colonies (Supplementary Fig. 2b) (n=3, human CD34+ donors, t-test *p-value< 0.01). Smaller colony sizes in *ALKBH5* gRNA-treated CD34^+^ cells could be attributed to a possible differentiation or proliferation block at the early progenitor stage of erythroid differentiation. Single-cell sequencing evaluation of the Methocult colonies was done in normal, and *ALKBH5* gRNA treated CD34^+^ cells (Supplementary Fig. 2c). As shown in supplementary Fig. 2c, the erythroid cluster was identified using expression of *GATA1*, *KLF1*, *HBB*, and *GYPA* mRNA expression while the myeloid cell cluster was marked by the *PTPRC* mRNA expression (Supplementary Fig. 2c). ALKBH5-deficient hCD34^+^ cells (number of cells analyzed n=5176) showed a clear differentiation block compared to the wildtype controls (number of cells analyzed n=5714 (Fig. 2c and supplementary Fig. 2d). Control gRNA-treated cells showed a singular trajectory denoted with a black line leading to the mature red cells, whereas the *ALKBH5* gRNA-treated cells showed a scrambled differentiation path. ALKBH5-deficient early erythroid precursors (red dots) showed more convoluted trajectory with disordered differentiating into mature red cells (marked with magenta dots) (Fig. 2c). This shows that. ALKBH5 is required for erythroid differentiation.

**Fig. 2.**
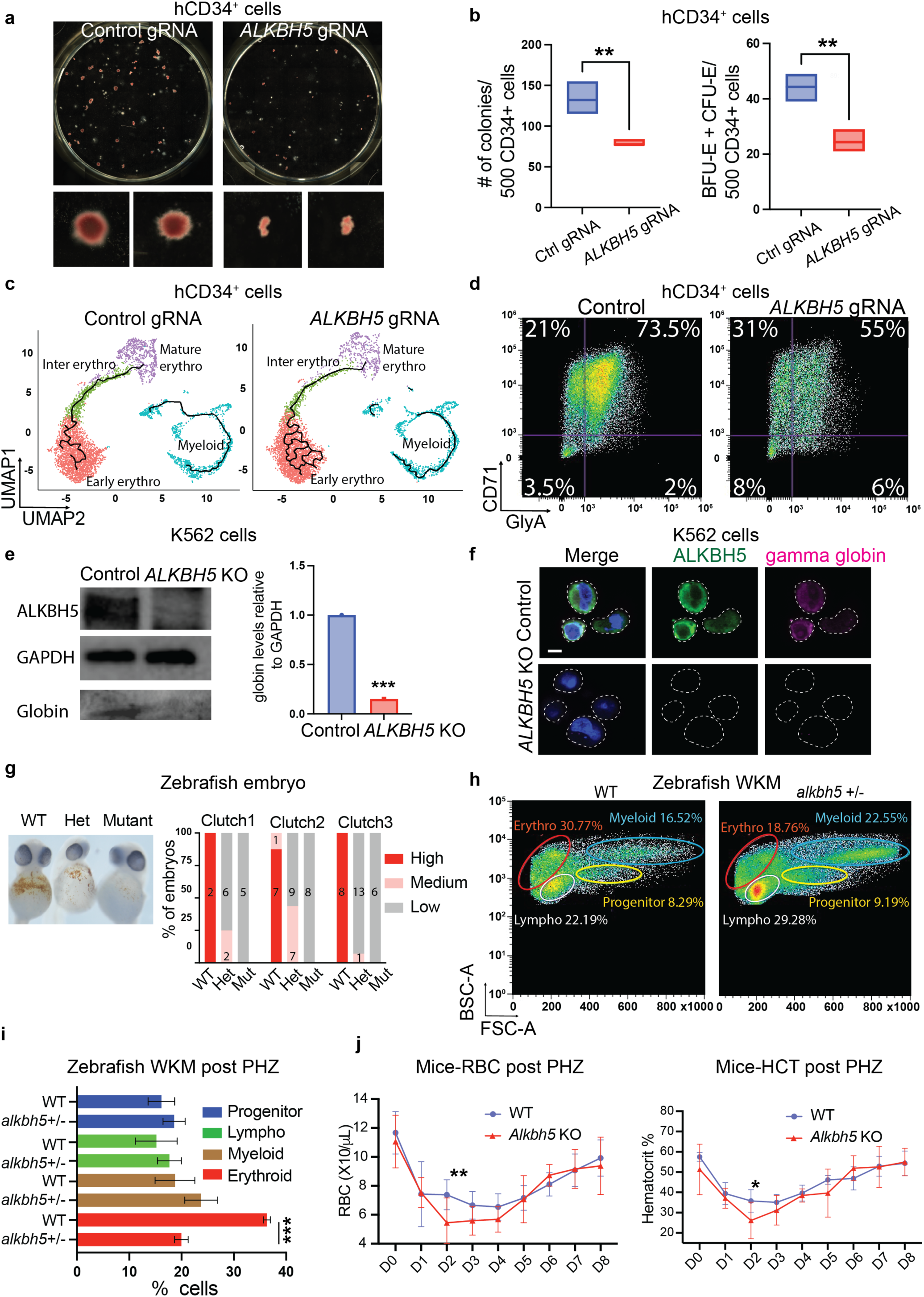
*ALKBH5* regulates erythropoiesis in human CD34+ cells and zebrafish, murine models. **a**, Methocult analysis of hCD34+ cells nucleofected with control gRNA or *ALKBH5* gRNA + Cas9. Representative colony from each plates depicting erythroid colony size in Control gRNA and *ALKBH5* gRNA. n=3 CD34+donors. **b**, Quantification of total and erythroid per 500 hCD34+ cells from methocult analysis. **c**, UMAP plots of control gRNA and *ALKBH5* gRNA hCD34+ cells with trajectory analysis. Red cluster shows early erythroid precursors, the green cluster represents intermediate erythroid precursors, and the magenta represents mature erythroid cells. Myeloid cells are shown in cyan color. **d**, FACS analysis of hCD34+ cells during erythropoiesis. Cells were transfected with Control gRNA or *ALKBH5* gRNA and subjected to FACS analysis on day7 using anti-CD71 and anti-CD235a antibodies. **e**, K562 *ALKBH5* KO cell analyzed for globin levels using western blot. K562 cells were probed on day 2 post hemin induction using anti-Alkbh5, anti-gamma globin and anti-GAPDH antibodies. **f**, Confocal microscopy of K562 *ALKBH5* KO cells using anti-ALKBH5, anti-gamma globin and Hoechst on day2 post hemin induction. **g**, Benzidine stain analysis of *alkbh5* +/− incross clutch larvae. Zebrafish larvae were subjected to Benzedine analysis and genotyping. The graph shows the quantification of the results from N=3 clutch. **h**, FACS analysis of Zebrafish whole kidney marrow (WKM). Adult zebrafish (6-12 month old) were processed for whole kidney marrow, and FACS was performed for blood cell counts. **i**, Graph shows different blood cell type distribution in WT and *alkbh5* +/− fish on day 8 of recovery of phenylhydrazine-induced acute anemia. **j**, Graph showing red blood cell recovery and hematocrit from day 0-8 in WT and Alkbh5 KO mouse model post-phenylhydrazine-induced anemia. n=7, WT mice, n=5, Alkbh5 KO mice. Scale bar-5 μm, p-values *** <0.001, ** <0.01 from unpaired t-test.

To further characterize this block in differentiation, we subjected ALKBH5 deficient hCD34^+^ and K562 cells to FACS analysis using antibodies against erythroid differentiation cell-surface markers, Transferrin receptor (CD71) and Glycophorin A (GlyA), (Choudhuri et al., 2020; Tamplin et al., 2015). Early erythropoiesis is marked by a transition from a CD71^+^GlyA^−^ to a CD71^+^GlyA^+^ state. The mature red cells are GlyA^+^ only (Choudhuri et al., 2020; Giarratana et al., 2011). Compared to the control, CD34^+^ cells with reduced ALKBH5 function failed to transition from the CD71^+^GlyA^−^ to the CD71^+^GlyA^+^ cells (Fig. 2d). We established an *ALKBH5*-KO K562 stable line harboring an early truncation mutation, predicted a complete loss of enzymatic activity (Supplementary Fig. 2e). Western blot analysis of the *ALKBH5* KO K562 line confirmed the absence of ALKBH5 protein (Fig. 2e and Supplementary Fig. 2e,f). The ALKBH5-KO K562 showed a reduction in globin production at two days after differentiation induction with hemin, probed using an anti-gamma-globin antibody (Fig. 2e). We validated these results using confocal microscopy of *ALKBH5* KO cells. Compared to control cells, *ALKBH5* KO cells showed the absence of ALKBH*5* protein in the cytoplasm and reduced levels of gamma globin (Fig. 2e, f). As shown in Supplementary Fig. 2g, we analyzed globin protein levels in control and mutant K562 cells using HPLC. The results showed a significant reduction in globin levels post-hemin induction in the mutant cells. In summary, hCD34^+^ stem and progenitor cells and K562 cells require *ALKBH5* function for erythropoiesis.

To examine function of alkbh5 in vivo, we generated an *alkbh5* KO Zebrafish mutant using CRISPR Cas9 targeting the *alkbh5* locus to confirm the gene function *in vivo* (Avagyan and Zon, 2016; Davidson and Zon, 2004). We targeted the *alkbh5* demethylase domain and introduced an early stop codon, thus rendering the protein non-functional (Supplementary Fig. 2h). Compared to wildtype and heterozygous sibling controls the *alkbh5* mutant F1 progeny showed a significant reduction in red cells assayed using the benzidine staining (Fig. 2g). Homozygous mutant embryos (N=3 clutches, n=19 embryos) showed low or no benzidine staining as shown in Fig. 2g, indicating embryonic anemia. *alkbh5* heterozygous embryos showed low red cells or anemia, signifying haploinsufficiency of *alkbh5* RNA demethylase function (n=38, 73%-low and 27%-medium benzidine stain). The wild-type siblings showed a normal benzidine level (n=18, 94%-high and 6%-medium benzidine stain). We observed low viability of *alkbh5* homozygous mutant fish and used *alkbh5* het fish for the FACS analysis of kidney marrow cells. The adult kidney marrow blood showed a significant reduction in red cells (Fig. 2h and supplement Fig. 2i-j) (Control-30.77%, *alkbh5* het 18.76%, **p-values<0.001), with an increase in the myeloid (Control-16.52%, *alkbh5*-het 22.55%, *p-values<0.01) and lymphoid compartments (Control-22.19%, *alkbh5*-het 29.28%) (Fig. 2h). Giemsa staining of the peripheral blood of *alkbh5* +/− zebrafish showed altered morphology of red blood cells with vacuolated cytoplasm (Supplementary Fig. 2i). We used phenylhydrazine (PHZ) induced acute anemia model to study red cell recovery in the *alkbh5* heterozygous mutant zebrafish. PHZ induces specific red cell membrane lysis and causes acute anemia (McReynolds et al., 2008). *alkbh5* Heterozygous and WT adult fish (age 4-12 months) were phenylhydrazine treated (0.003mg/ml) for 20 mins and followed for red cell recovery and analyzed by FACS on day 8 post-PHZ treatment as described by McReynolds et al. (McReynolds et al., 2008). As shown in Fig. 2i and supplementary Fig. 2k, *alkbh5* het zebrafish failed to recover red cells (n=6, Control-36.2%, n=10, *alkbh5* het 19.5%, ***p-values<0.0001) by the whole kidney marrow (WKM) blood analysis. The response to anemic stress was deficient in the *alkbh5* heterozygous mutants.

*Alkbh5* depletion in mice delayed the development and progression of leukemia induced by oncogenes (Cheng et al., 2020; Shen et al., 2020; Wang et al., 2020). Mice *Alkbh5* mutations accelerate leukemia, but steady-state hematopoiesis is relatively normal. We hypothesized that murine *Alkbh5* KO erythroid progenitors would have difficulty responding to acute stress, as demonstrated in the case of zebrafish models. Similar to zebrafish, we used a non-lethal single dose of PHZ (100 mg/kg mouse) to lyse 50% of erythrocytes inducing anemia. WT C57BL/6 mice fully recover hematocrits in 5–7 days post PHZ treatments. We then compared PHZ-treated *Alkbh5*^−/−^ to *Alkbh5*^+/+−^ littermates for their hematocrit recovery. As shown in Fig. 2j Alkbh5^−/−^ mouse RBC counts (5.28 ×10^6^/μL compared to *Alkbh5*^+/+−^ mice-7.38 ×10^6^/μL, p-value-0.023 t-test) and hematocrit were significantly lower (26.14% compared to WT-35.76%, p-value-0.034, t-test) on D2-D3 post-PHZ, indicating *Alkbh5* KO mice were defective in stress erythropoiesis. In summary, stress-induced acute anemia in *Alkbh5^−/−^*mice showed defective erythrocyte recovery, indicating its role in blood regeneration under stress. Our examination of erythropoiesis of mutant hCD34^+^ and K562 cells *in vitro* and the zebrafish and murine KO model *in vivo* established the required role of *ALKBH5* RNA demethylase in stress erythropoiesis.

### Stress granule core component mRNAs are m^6^A modified

ALKBH5 is a key RNA demethylase that directly controls the transcriptome m^6^A levels (Mauer and Jaffrey, 2018; Zhang et al., 2017; Zheng and He, 2015). To validate and accurately map m^6^A marks on transcripts targeted by ALKBH5 activity, we performed anti-m^6^A antibody-independent m^6^A transcriptome profiling by the MazF digestion of RNAs isolated from hCD34+ cells after Alkbh5 CRISPR-Cas9 editing (Protocol details in the method material section). In this approach, RNA endonuclease, MazF, specifically cleaves single-stranded RNAs at the 5’ side immediate to unmethylated (ACA) sequence, but not methylated (m^6^ACA). Total RNA without MazF digestion serves as input control for the comparison with MazF-cleaved RNA fragments. Each fraction is then converted into cRNA (Complementary RNA) with unique fluorescent labels and hybridized with m6A Single Nucleotide Array probes. 14,321 unique probes (60 bp) that identify 11,237 m^6^A sites, giving the quantitative value on the m^6^A level at single nucleotide resolution. m^6^A Microarray does not use m^6^A-antibody immunoprecipitation, thus validating our meRIP-seq results independently. m^6^A analysis of *ALKBH5* KO CD34+ cells during erythropoiesis showed m^6^A upregulation of 4800 genes in comparison to control gRNA cells (Supplementary Fig. 3a). Pathway analysis using all the m^6^A marked transcripts revealed Spinocerebellar Ataxia-2 pathway upregulation with *ALKBH5* gRNA perturbation (Fig 3a) (Table 2). m^6^A levels of various stress granules and RNA metabolism genes were significantly upregulated in *ALKBH5* KO CD34^+^ cells. We found a specific m^6^A signature in 3’UTR of *ATXN2*, *G3BP*, and *Tia1* genes (Fig. 3b). We validated ALKBH5 mediated ATXN2 regulation in *ALKBH5* KO K562 model using MeRIP sequencing (Fig. 3c). Quantification of the m^6^A levels of *ATXN2* mRNA showed an elevation in *ALKBH5* KO cells compared to the control cells (Fig. 3c, d). m^6^A abundance analysis of erythroid-specific genes such as *HBA*, and *GATA1* showed an increase in m^6^A levels in mRNA 3’UTR regions (Fig. 3d). RNA binding proteins such as *G3BP, TIA1, YTHDF2, HNRNPA1* revealed mRNA 3’ UTR m^6^A upregulation. GAPDH mRNA exhibited no such changes in m^6^A levels for *ALKBH5* KO cells (Fig. 3d). Quantification of m^6^A on *ATXN2* transcripts indicated a 14-fold 3’ UTR m^6^A upregulation regulation (Fig. 3d, **p-value< 0.001). These results demonstrate m^6^A RNA modification dynamics of SG core component genes and validate their regulation via ALKBH*5* demethylase activity.

**Fig. 3.**
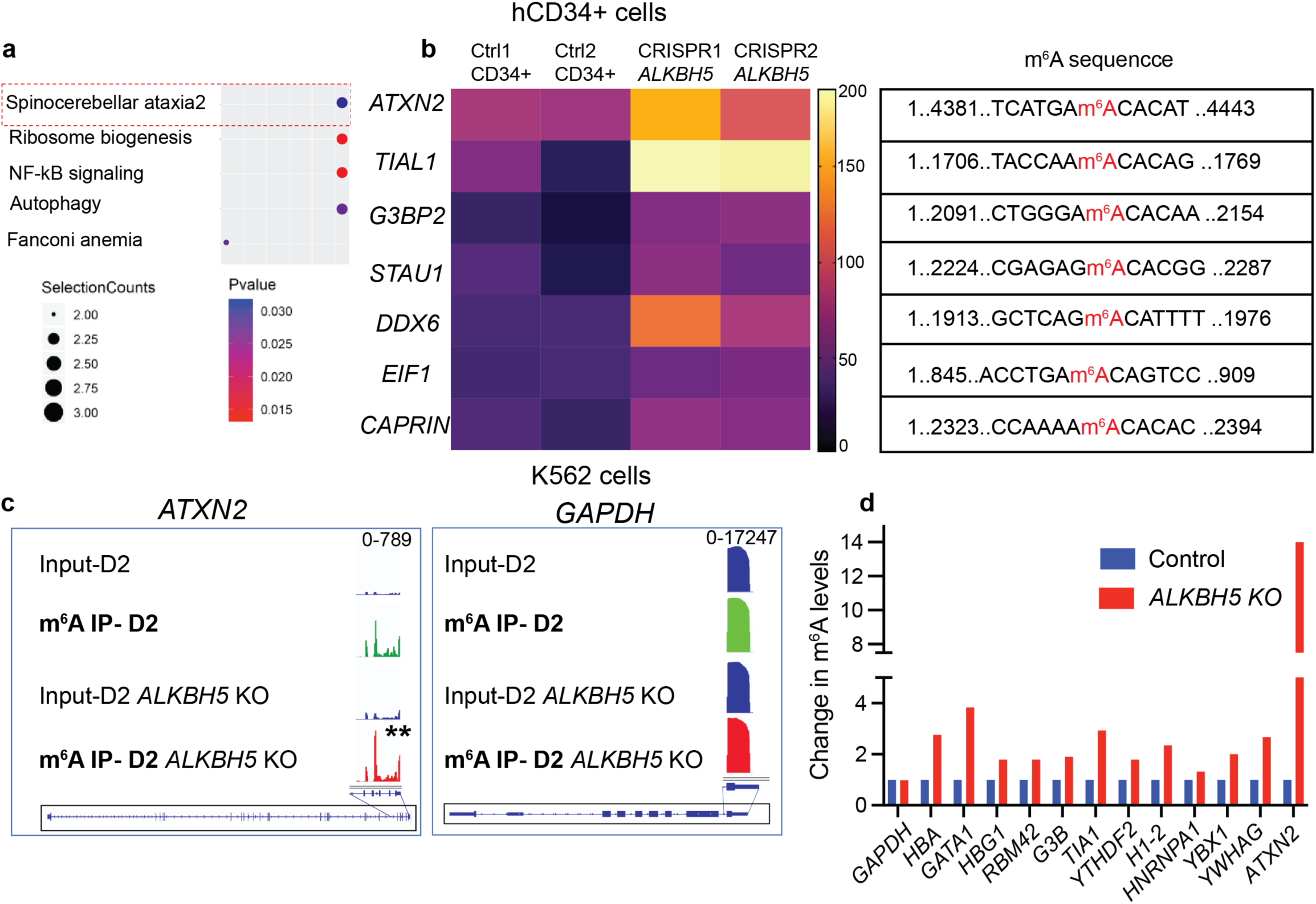
ALKBH5 regulates SG core gene demethylation during erythropoiesis. **a**, Pathway analysis of transcriptome-wide m^6^A analysis from *ALKBH5* gRNA and control gRNA hCD34+ cells (day 7). Pathways such as spinocerebellar ataxia2, ribosome biogenesis, and other cellular signaling are upregulated in *ALKBH5* gRNA-treated cells. **b**, m^6^A analysis showing significant upregulation of methylation on SG core gene in *ALKBH5* gRNA cells. *ATXN2*, *TIAL1*, *G3BP2*, *STAU1*, and other RBPs show high 3’UTR m^6^A levels in *ALKBH5* gRNA hCD34+ cells. The table depicts the 3’UTR sequence with the site of methylation. **c**, m^6^A levels on *ATXN2* mRNA in *ALKBH5* KO K562 cells during hemin-induced differentiation and for control gene GAPDH. **d**, Graph showing m^6^A levels of key erythropoiesis and RNA metabolism genes in K562 *ALKBH5* KO cells day 2 post hemin induction.

**Table 2.**
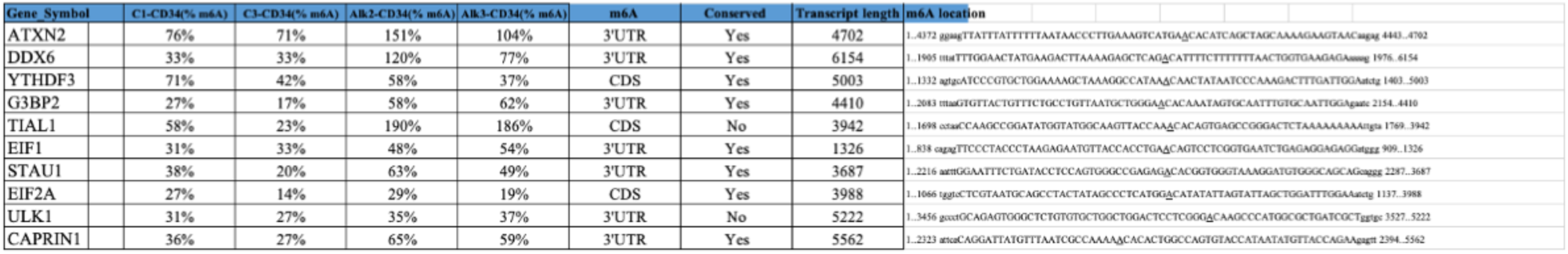
m^6^A microarray analysis of hCD34^+^ ALKBH5 KO and control gRNA treated cells.

### ATXN2 RNA m^6^A impact stress granule assembly and regulate erythropoiesis

mRNA m^6^A levels are reported to impact the protein translation (Zaccara et al., 2019; Zheng and He, 2015). To decipher the mechanism underlying the control of ALKBH5-mediated m^6^A levels on cell proteome and levels of various SG core components, we performed mass spectrometry on hCD34^+^ cells. Our proteomic analysis of *ALKBH5* KO and control erythroid cells showed a significant downregulation of diverse essential genes for mRNA translation, RNA metabolism, and stress granule biology (Fig. 4a). ATXN2 (Spinocerebellar ataxia 2) is one of the proteins that showed significant downregulation. ATXN2 is crucial in translation regulation and directly impacts cell proteome via SG-mediated ribostasis (Lastres-Becker et al., 2016; Ramaswami et al., 2013). We validated these findings using *ALKBH5* KO K562 cells and showed significant downregulation of ATXN2 protein levels in comparison to control K562 cells (Supplementary Fig. 4a). The role of ATXN2 and SGs in blood development is novel and largely remains elusive. G3BP-mediated SG assembly is key for coordinating the production of α-globin and heme, while ATXN2 has been shown to regulate megakaryopoiesis via timing translation of surface receptor ITGB3 and PECAM1 (Ghisolfi et al., 2012; Hansen et al., 2020). hCD34^+^ cells with *ATXN2* gRNA showed a significant decrease in CFU-E and BFU-E formation capacity of hCD34^+^ cells in colony-forming assays. (Fig. 4b, n= 2 human CD34^+^ cell donors, with 3 technical replicates. **P-value<0.001). In summary, ALKBH5 controls ATXN2 protein levels required for hCD34+ erythroid differentiation.

**Fig. 4.**
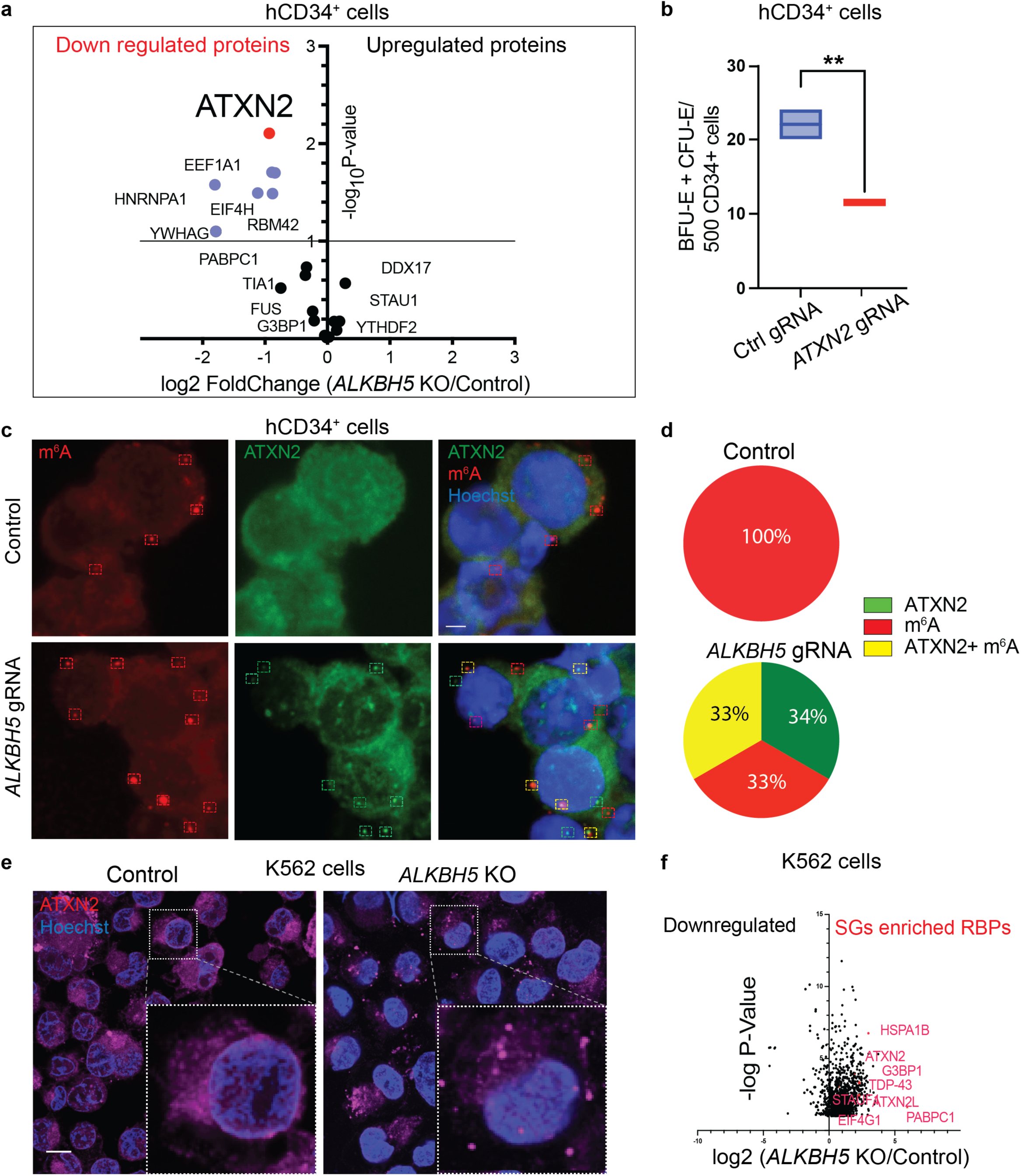
*ATXN2* RNA m^6^A levels impact stress granule assembly and regulate erythropoiesis. **a**, Mass spectrometry analysis of the total proteome of control gRNA and *ALKBH5* KO hCD34+ cells. The left side of the graph shows downregulated and the right side indicates upregulated protein in *ALKBH5* KO cells. **b**, Methocult analysis of hCD34+ control and *ATXN2* KO cells. Quantification of BFU-E/CFU-E per 500 hCD34+ cells for control gRNA and *ATXN2* gRNA treated hCD34+ cells n=2 hCD34+ donors. **c**, STED microscopy of hCD34+ cells during erythropoiesis. hCD34+ cells were subjected to control gRNA or *ALKBH5* gRNA. Cells were probed using anti-m^6^A, anti-ATXN2 antibodies, and Hoechst. Dotted rectangles show puncta labeled with anti-m^6^A (red rectangles), anti-ATXN2 (green rectangle), and anti-m^6^A + anti-ATXN2 antibodies (yellow rectangles). **d**, Quantification of anti-ATXN2, anti-m^6^A, ATXN2+ m^6^A labelled puncta from control and *ALKBH5* KO hCD34^+^ cells. n=3, N=20 cells. **e**, STED microscopy of control and *ALKBH5* KO K562 cells stained with anti-ATXN2 and Hoechst. n=3. **f**, anti-ATXN2 antibody pulldown and Mass spectrometry analysis of proteins from control gRNA and *ALKBH5* KO K562 cells. scale bar-5 μm, p-values *** <0.001, ** <0.01 from unpaired t-test.

To confirm *ATXN2’s* role in erythropoiesis, we used siRNA-mediated knockdown combined with western blot analysis in hemin-induced K562 cells. Downregulation of ATXN2 protein during erythropoiesis abrogated gamma-globin production (Supplementary Fig. 4b). Anti-GAPDH antibody staining showed equal levels for control siRNA-treated K562 cells and *ATXN2* siRNA-treated cells (Supplementary Fig. 4b). We confirmed this result using benzidine staining (an indicator of globin levels) of control siRNA-treated and *ATXN2* siRNA-treated K562 cells. As shown in (Supplementary Fig. 4c), ATXN2 downregulation significantly decreases benzidine stain, denoting failure of erythropoiesis and leading to anemia-like symptoms.

To confirm these results, we CRISPR-edited *ATXN2* in K562 cells and subjected them to hemin induction. FACS analysis of *ATXN2* gRNA cells using anti-CD71 and anti-GlyA surface marker antibodies showed a clear block in cell differentiation compared to control cells (Supplementary Fig. 4d). Hemin-induced control K562 cells transitioned from CD71^+^, GlyA^−^ (48.8%) to CD71+, GlyA+ cells (34.5%). *ATXN2* gRNA cells failed to transition from CD71+, GlyA- (59.1%) to CD71^+^, GlyA^+^ cells (13.2%), indicating an early erythropoiesis defect (Supplementary Fig. 4d). Interestingly, almost twice (CD71^−^ GlyA^−^, 27% of total cells) *ATXN2* gRNA K562+ cells showed a clear failure to take up erythroid fate compared to control cells (CD71^−^ GlyA^−^, 14.2%). To obtain insight into ATXN2 levels in *ALKBH5* KO models, we used an anti-ATXN2 antibody and performed super-resolution microscopy. We probed *ALKBH5* CRISPR cells during hCD34^+^ cell erythropoiesis using an anti-ATXN2 antibody and anti-m^6^A antibody (Fig. 4c). *ALKBH5* CRISPR cells showed ATXN2 SGs with numerous puncta co-labeled with anti-m^6^A antibody. We also observed anti-ATXN2 only and anti-m^6^A only puncta in the cells (Fig. 4c, d). The absence or reduction of *ALKBH5* RNA demethylase results in altered ATXN2 SGs that could sequester m^6^A-modified RNAs and other key SG RBPs.

To study the SGs in further detail, we confirmed these SGs results using K562 *ALKBH5* KO models. We used an anti-ATXN2 antibody and performed super-resolution microscopy during hemin-induced erythropoiesis. Control K562 cells showed an even cytoplasmic distribution of ATXN2 protein (Fig. 4e). *ALKBH5* KO cells showed massive SGs in the cytoplasm (Fig. 4e). To characterize SGs in Alkbh5 KO models, we performed anti-ATXN2 antibody-mediated pulldown and mass spectrometry analysis. In comparison to control cells, anti-ATXN2 antibody pulldown from *ALKBH5* KO cells showed many fold enrichment of various known proteins of SGs, such as TDP-43, G3BP1/2, PABP, Tia1, and other RBPs enriched in SGs (Yang et al., 2020)(Fig. 4f) (−logP-value>2). We investigated some of these SGs markers during erythropoiesis. SGs are reported to sequester RNAs subjected to various fates based on the cell state and various physiological conditions (Anderson and Kedersha, 2009; Ramaswami et al., 2013; Tauber et al., 2020). We used an anti-m^6^A antibody and an anti-G3BP antibody to mark RNAs in SGs. As shown in supplementary Fig. 4e, hCD34^+^ cells with *ALKBH5* gRNA CRISPR revealed SGs labeled with m^6^A RNAs. In control, gRNA CRISPR hCD34^+^ cells, m^6^A labeled RNAs were evenly distributed, and G3BP distribution was unaffected (supplementary Fig. 4e). Our combined results from *ALKBH5* KO K562 and hCD34^+^ cell whole-cell proteome mass spectrometry, super-resolution microscopy, and pulldown of SG proteins correspond to SG-mediated m^6^A mRNA accumulation.

### SGs core component RNA m^6^A levels regulate active translation

To assess a ribosome-mediated translation regulation of mRNAs with m^6^A-modified 3’UTR during erythropoiesis, we performed hCD34^+^ cells polysome analysis during erythropoiesis. hCD34^+^ cells were knocked out for the *ALKBH5* gene and subjected to polysome analysis using a Sucrose gradient (5-50%) (Fig. 5a). Control cell polysome trace is shown in green, while the red graph represents *ALKBH5* gRNA cells (Fig. 5a). As predicted, *ALKBH5* KO cells hCD34+ cells showed a marked reduction in actively translating heavy polysomes (Fig. 5a). Our super-resolution microscopy results shown in Fig. 4c, d corresponded to the enrichment of m^6^A mRNAs in a non-translating pool represented by the pre-polysome fraction in *ALKBH5* KO cells (Fig. 5a). Non-translating mRNAs are reported to localize to SGs (Balagopal and Parker, 2009; Chantarachot and Bailey-Serres, 2018). We subjected the pre-polysome fraction marked by a dotted box to RNA sequencing (Fig. 5a) and comparative analysis to identify and measure different mRNA populations. As shown in the volcano plot (Fig. 5b), it revealed a significant enrichment of various mRNAs in the light non-translating pre-ribosomal pool in *ALKBH5* KO hCD34^+^ cells (Fig. 5b). As predicted, *ALKBH5* gRNA treated hCD34^+^ cells showed mRNAs for *ATXN2*, *TIAL1*, *EIF1*, *CAPRIN1*, and other SGs core components enriched in non-translating fraction. Our single nucleotide m^6^A analysis previously revealed a significant upregulation of methylation on these transcripts in the *ALKBH5* gRNA treated hCD34+ cells (Fig. 3b). To confirm mRNA m^6^A 3’UTR mediated translation regulation, we used 3’UTR GFP reporter constructs. We transfected K562 cells with a control 3’ UTR or the *ATXN2* gene 3’UTR tagged to the GFP to validate the effect of ALKBH5 on erythropoiesis. Control 3’UTR sequence contained no consensus m^6^A motif GGAC identified in our analysis (Fig. 1e), but *ATXN2* gene 3’UTR contains the sequence that showed m^6^A modification identified in K562 and hCD34^+^ cell m^6^A analysis (Fig. 3). Confocal microscopy of hemin-induced K562 cells was performed for control 3’ UTR-GFP and *ATXN2* 3’UTR GFP. Control 3’ UTR GFP levels remained the same and were unaffected in both WT K562 and *ALKBH5* KO cells, but *ATXN2* 3’UTR GFP reporter levels were significantly downregulated in *ALKBH5* KO cells (Fig. 5c). This demonstrates that *ALKBH5* RNA demethylase activity is crucial in regulating protein levels via 3’UTR m^6^A levels. *ATXN2* mRNA and other SGs mRNA are translationally blocked in *ALKBH5* KO cells. In summary, high m^6^A modification at 3’UTR of SG core gene *ATXN2* mRNAs in *ALKBH5* KO cells reduces protein levels. Decreased ATXN2 leads to SG formation that sequesters mRNAs decreasing global translation and leading to are reduction of erythroid cells.

**Fig. 5.**
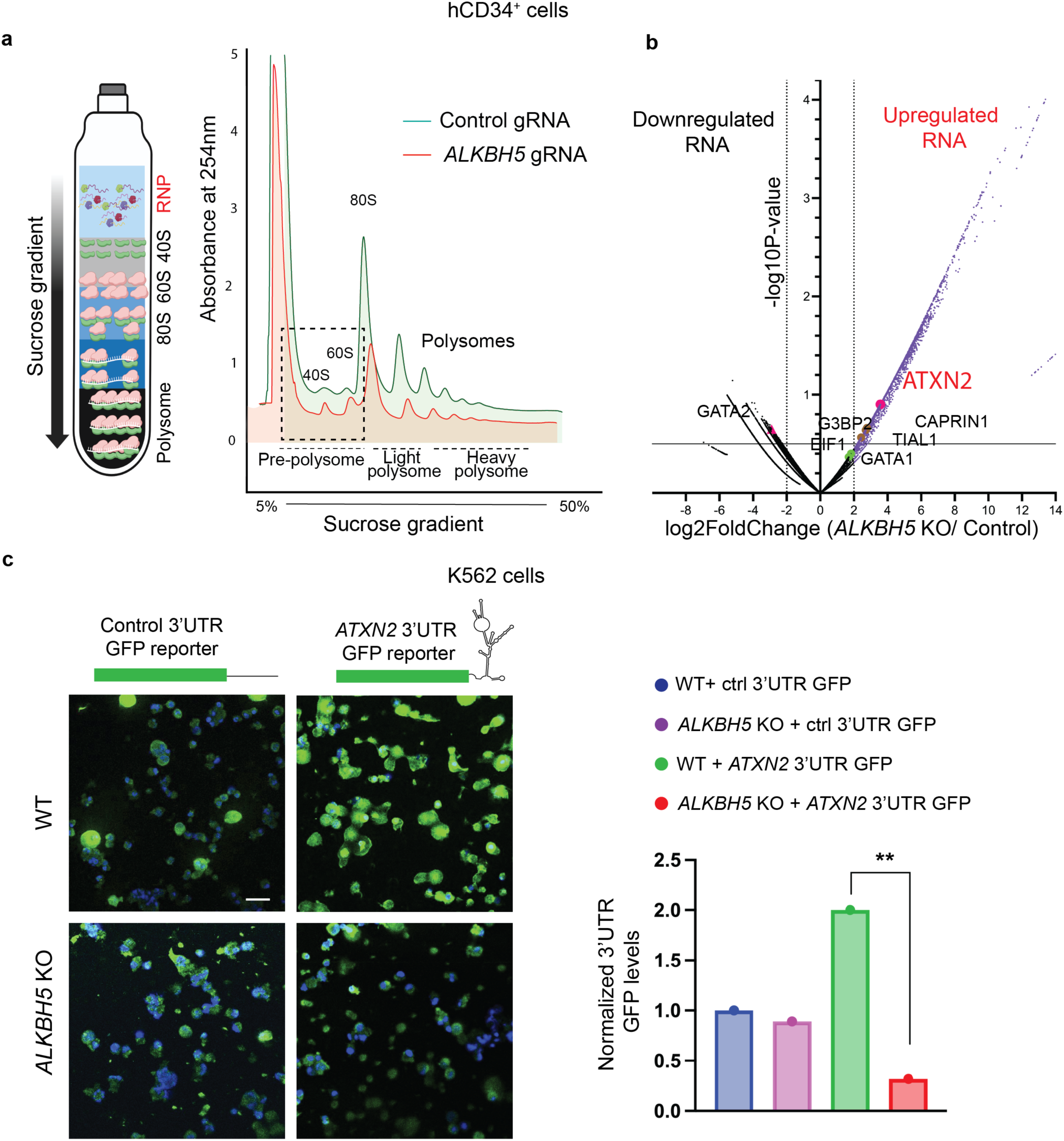
3’UTR mediated translation control mechanism regulates SG protein levels. **a**, Polysome analysis of hCD34+ control and *ALKBH5* KO cells. Cells were subjected to CRISPR editing using control gRNA or *ALKBH5* gRNA and processed for polysome analysis during erythropoiesis (Day 4). **b**, Volcano plot of RNA sequencing of the pre-polysome fraction of control and *ALKBH5* gRNA hCD34+ cell. *ATXN2* RNA and *G3BP2, TIAL1, CAPRIN1* coding for SG core components enriched in the non-translating pool in *ALKBH5* KO hCD34+ cells. **c**, K562 WT and *ALKBH5* KO cells transfected with GFP reporter tagged to control 3’UTR sequence and *ATXN2* gene 3’UTR sequence. Post-transfection cells were hemin induced, and GFP expression (green) levels were analyzed using the confocal microscope. Cells were co-stained with DAPI (blue) n=3, N=30 cells Scale bar-10μm, p-values ** <0.01 from unpaired t-test.

### ATXN2-mediated SG assembly regulates human erythropoiesis

ATXN2 is crucial in ribostasis via stress granule regulation, and mutations are known to cause spinocerebellar ataxia-2 in humans (Elden et al., 2010; Lastres-Becker et al., 2016; Ramaswami et al., 2013). To validate the role of ATXN2 protein in blood development and homeostasis, we overexpressed *ATXN2* in hCD34^+^ cells and measured its effect on erythropoiesis. Lentiviral-mediated *ATXN2* overexpression (*ATXN2* OE) in hCD34^+^ cells enhanced erythroid differentiation compared to GFP controls as early as day 2 of differentiation culture (Fig 6a). FACS analysis for surface markers using anti-CD71 and anti-GlyA antibodies of *ATXN2* OE hCD34^+^ cells showed an early transition from CD71^+^, GlyA^−^ to CD71^+^, GlyA^+^ cells (30.1%) in comparison to control cells (2%). We subjected *ATXN2* OE hCD34^+^ cells to a Methocult colony formation assay and measured the number of erythroid colonies. As shown in supplementary Fig. 5a, *ATXN2* OE led to larger and supernumerary erythroid colonies. We confirmed the pro-erythroid fate effect of *ATXN2* overexpression using K562 cells. K562 cells were transfected with either GFP lenti viral vector or an *ATXN2* overexpression vector and assayed using anti-CD71 and anti-GlyA surface marker antibodies during hemin-induced erythropoiesis (Supplementary Fig. 5b). ATXN2 overexpression led to a significantly high number of CD71^+^, GlyA^+^ cells in *ATXN2* overexpression(71%) in comparison to control cells (59.9%). *ATXN2* expression promoted a significantly higher number of cells transitioning from early CD71^+^, GlyA^−^ state to CD71^+^, GlyA^+^ state. In summary, these results demonstrate that *ATXN2* promotes erythroid differentiation.

**Fig. 6.**
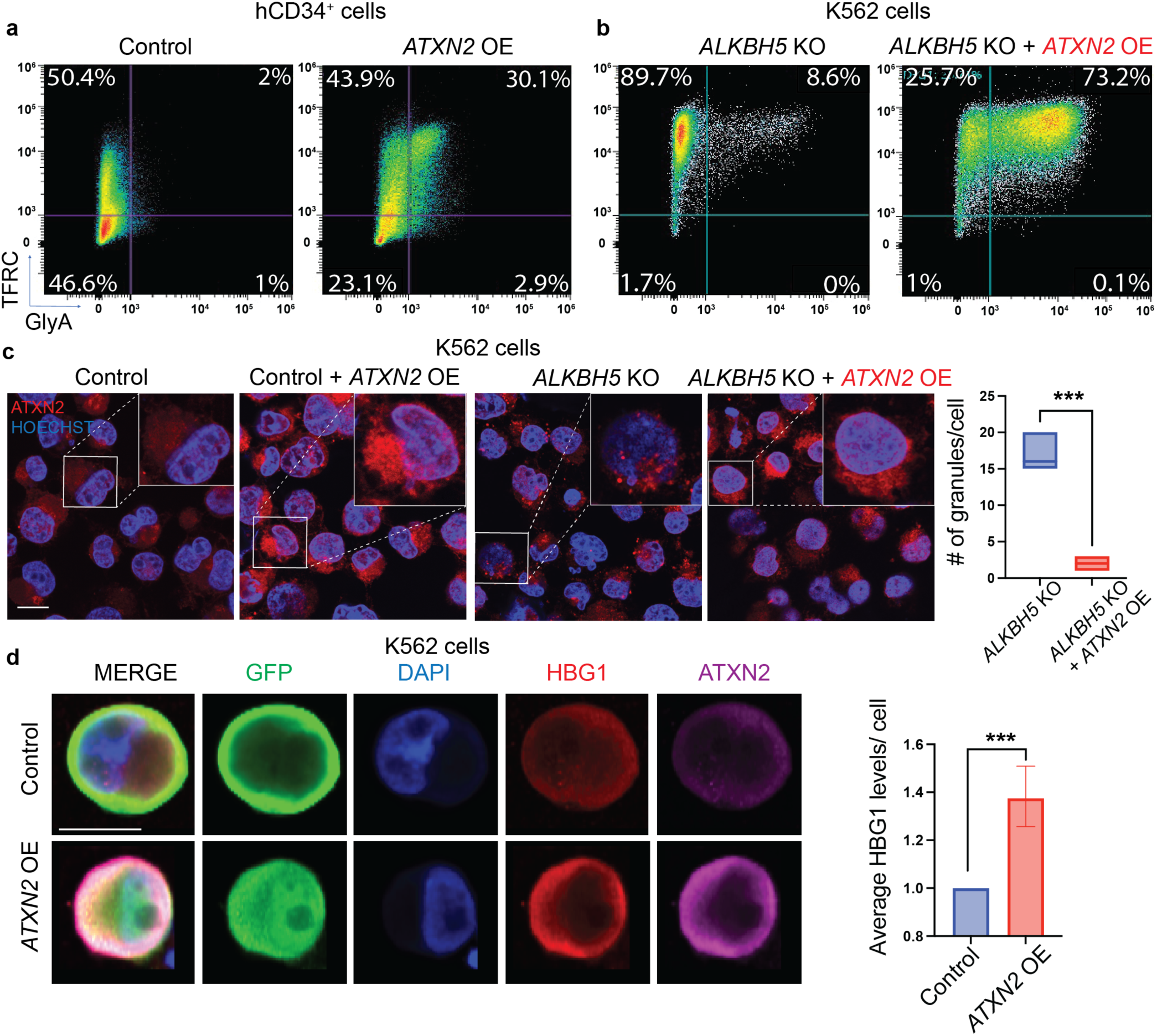
ATXN2 promotes and rescues erythropoiesis. **a**, FACS analysis of CD34+ cells during erythropoiesis. Cells were transfected with Control lenti vector or ATXN2 overexpression (OE) lenti vector and subjected to FACS analysis on differentiation day 2 using anti-CD71 and anti-CD235a. n=2 independent human CD34+ donors **b**, FACS analysis of K562 Alkbh5 KO cells transfected with Control lenti vector or ATXN2 OE lenti vector. Cells were subjected FACS analysis using anti-CD71 and anti-CD235a on day 2 post hemin induction. **c**, STED microscopy of Control, Alkbh5 KO, control + ATXN2 OE and Alkbh5 KO+ ATXN2 OE K562 cells stained using anti-ATXN2 antibody transfected with control or ATXN2 OE lenti vectors on day2 hemin induced erythropoiesis. Graph showing number of stress granules in Alkbh5 KO and Alkbh5 KO + ATXN2 OE K562 cells. **d**, K562 cell probed for HBG1 and ATXN2 during hemin induction. K562 cells were transfected with control or ATXN2 overexpression lenti vector. Cells were stained using anti-GFP (green), DAPI (blue), anti-gamma globin (red) and anti-ATXN2 (magenta). Quantification of control and ATXN2 OE K562 cells stained using an anti-HBG1 antibody. n=4 experiments, N=20 cells for quantification, Scale bar-5 μm, p-values *** <0.001, ** <0.01 from unpaired t-test.

To study *ATNX2* and stress granule formation in depth, we overexpressed *ATXN2* in *ALKBH5* KO K562 cells. Our *ALKBH5* KO results showed an erythroid differentiation block (Fig. 6b and Fig. 2). *ALKBH5* downregulation reduced ATXN2 protein levels (Fig. 4A). We replenished ATXN2 in *ALKBH5* KO cells using lentiviral *ATXN2* overexpression. Restoring ATXN2 levels in *ALKBH5* KO cells led to the rescue of K562 cell erythropoiesis assayed using FACS analysis of cells for CD71, GlyA surface markers during d2 post hemin induction (Fig. 6b). *ATXN2* overexpression in *ALKBH5* KO promoted cells to CD71^+^, GlyA^+^ (73.2%) substantially removing the differentiation block seen in the case of *ALKBH5* KO cells (8.6%) (***p-value< 0.0001). To delineate the cell biology of SGs, we combined super-resolution microscopy with the anti-ATXN2 antibody staining in K562 control, *ATXN2* overexpressing WT K562, *ALKBH5* KO, and *ATXN2* overexpressing *ALKBH5* KO K562 cells. As shown previously, *ALKBH5* KO cells showed large SGs (Fig. 6c). *ATXN2* overexpression in *ALKBH5* KO cells ultimately resolved the SG phenotype and showed a normal cytoplasmic distribution of ATXN2 protein (Fig. 6c, ***p-value< 0.0001). *ATXN2* overexpression did not change the phenotype of WT K562 cells (Fig. 6c). Immunostaining using anti-PABP protein antibody combined with super-resolution microscopy revealed an increase in the presence of PABP protein SGs colocalizing with m^6^A RNA probed using the anti-m^6^A antibody in *ALKBH5* KO cells in comparison to control K562 during erythroid differentiation (Supplementary Fig. 5c, **p-value< 0.001). ATXN2 overexpression in *ALKBH5* KO cells led to complete resolution of SGs, while *ATXN2* overexpression in control K562 cells showed no change in PABP and m^6^A RNA distribution (Supplementary Fig. 5c). To validate the ATXN2 role in erythropoiesis we probed for gamma-globin using confocal microscopy. We used *ATXN2* over-expressing K562 cells to probe for gamma-globin post-hemin induction. Control cells were similarly transfected with the control lenti vector and subjected to hemin induction. Confocal microscopy done using an anti-ATXN2 antibody showed ATXN2 upregulation in overexpression constructs in comparison to control K562 cells (Fig. 6d). Gamma globin measurements with anti-gamma globin antibody similarly showed up-regulation in comparison to control K562 cells (Fig. 6d) (***p-values < 0.0001). In conclusion, ATXN2 regulates SG assembly and functions as a pro-erythroid differentiation factor promoting globin protein production and erythroid cell differentiation.

## Discussion

Our study highlights a direct link between RNA m^6^A modification and the regulation of stress granule assembly during erythropoiesis. Numerous RNAs are transcribed during erythropoiesis and are m^6^A-modified. High m^6^A levels in RNAs keep these RNAs in the translationally suppressed state. Stage-specific RNA demethylation and concomitant translation are key to erythropoiesis. Erythroid and SG core protein mRNAs are m^6^A-modified and demethylated during differentiation by ALKBH5 RNA demethylase. *ALKBH5* mutant hCD34+ cells show significantly decreased levels of SG proteins, such as ATXN2, promoting SG formation. SGs accumulate mRNAs inhibiting the translation of numerous m^6^A-modified RNAs, causing erythropoiesis failure. Restoring the *ATXN2* function rescued the erythroid differentiation of *ALKBH5* mutant cells and reduced SG numbers to normal numbers in *ALKBH5* mutants. In this case, mRNAs released from SGs are translated by ribosomes, thus accelerating erythroid differentiation during stress. In the *ALKBH5* mutant erythroid cells, m^6^A-modified RNAs accumulate in the newly formed SGs marked by RBPs such as G3BP and PABP. *ALKBH5* mutant hCD34+ cell polysome experiments showed SG-specific mRNAs in the non-translating pool. *ATXN2* overexpression in normal hCD34+ results in enhanced erythropoiesis, with significantly increased erythroid colony formation demonstrating pro-translation function. ATXN2 is reported to facilitate RNA loading onto functional ribosomes (Lastres-Becker et al., 2016). This work establishes a mechanism by which ATXN2 facilitates the better translation of mRNAs in normal hCD34+ erythropoiesis and the release of SG-stored m^6^A-modified RNAs during stress-induced erythroid differentiation.

*ALKBH5* has been studied in murine blood cells, but there is no apparent hematopoietic phenotype at a steady state. These mice are prone to leukemia in response to oncogenes (Shen et al., 2018; Zheng et al., 2013). In contrast to the mice, *alkbh5* homozygous zebrafish mutants have anemia and *alkbh5* heterozygous mutants fail to regenerate red blood cells after phenyl hydrazine-induced stress. *ALKBH5* mutant hCD34+ cells have a block in erythroid differentiation. Stress conditions in our zebrafish and mice *ALKBH5* KO leads to the failure of erythropoiesis, causing anemia. Embryonic erythropoiesis is considered a stressful situation as the organism attempts to rapidly differentiate mesoderm into red blood cells to meet the oxygen requirement (Bertrand and Traver, 2009; Davidson and Zon, 2004; Tan et al., 2016). The *ALKBH5* mouse mutant also has a spermatogenesis defect, where rapid production of sperm is subject to stress. In the culture of human blood cells, cytokines and conditions create stress. Our results suggest that stress conditions during erythropoiesis (Paulson et al., 2011; Perry et al., 2009) accelerate a phenotype in the *ALKBH5* mutant cells.

m^6^A-modified RNAs are known to localize to SGs (Fu and Zhuang, 2020), and our work suggests that the RNAs are sequestered in SGs and are untranslated. m^6^A labeled mRNAs from SGs are reported to be released post-stress relief and translated in U2OS and HEK-TIA1 cells following sodium arsenite-induced stress (Das et al., 2022), although the mechanism remains unknown. Super-resolution microscopy demonstrated the presence of m^6^A-labeled RNA in the SGs showing their role as a cytoplasmic storage depot for RNA (Fig. 4N-P). A large number of erythroid-specific and SG-specific m^6^A modified RNAs were detected with MeRIP seq, and these would likely be present in SGs of erythroid progenitors. Release of these RNAs from stress granules to ribosomes would enhance erythropoiesis by allowing immediate translation of erythroid-specific mRNAs.

ATXN2 promotes the loading of cell-specific RNAs onto functional ribosomes(Lastres-Becker et al., 2016; Li et al., 2022). Overexpression of *ATXN2* leads to enhanced differentiation of erythropoiesis in normal human CD34+ cells. *ATNX2* overexpression also reduces SG number in *ALKBH5* mutant cells and rescues their differentiation. Our anti-ATXN2 antibody pulldown, mass spectrometry data, and microscopy ultimately showed the enrichment of various RNA-binding proteins in SGs. Given that ATXN2 itself is a crucial protein of SGs, the overexpression must alter the stoichiometry of the SG core complex, facilitating the dissolution of SGs and the loading of RNAs onto ribosomes. ATXN2 binds to PABP and 48S pre-initiation components in neurons and mouse embryonic fibroblasts to enhance the mRNA translation (Lastres-Becker et al., 2016). *ATXN2* overexpression rescues poly-Q mutant ATXN-3 mediated translation defects in the Machado-Joseph neurodegenerative disease models (Nóbrega et al., 2015). We identified ATXN2 as a pro-translation factor during erythropoiesis during stress. Our work suggests that ALKBH5 demethylase function is also crucial in regulating the stoichiometry of SGs core components via 3’UTR m^6^A. In summary, ATXN2 functions as a SG structural protein but can also resolve stress granules, allowing specific RNAs regulating differentiation to be loaded onto ribosomes.

CAG expansion and neuropathies caused by SG disease mutations lead to altered ribostasis affecting the protein synthesis (Elden et al., 2010; Sellier et al., 2016; Sproviero et al., 2017). *ATXN2* mutations are known to affect different age groups based on the number of repeat expansions (Bakthavachalu et al., 2018; Lastres-Becker et al., 2016; Singh et al., 2021). We recently examined blood counts from a recent clinical trial by Biogen including patients with CAG expansions in various SG protein genes, including *ATXN2* mutant patients. Human CAG poly-Q expansion of intermediate length showed no red blood cell phenotypes in Human patients (Age ~50-55 years) observed for 18 months. No bone marrow evaluation was done, and SGs in hematopoietic lineages were not studied. This data show that diseased poly-Q *ATXN2* does not lead to anemia in homeostasis conditions.

The level of m^6^A-modified RNA regulates erythropoiesis via translational control. SG mRNA m^6^A levels control core protein levels and thus regulate SG formation. SGs are formed during cellular stress conditions and regulate ribostasis. Our findings point to SG core component roles in possible disease-related complications, i.e., anemia and other blood disorders (Choudhuri et al., 2020). Acute anemia, thalassemia, and sickle cell disorders represent stressed hematopoiesis conditions. Our findings suggest a crucial role of SGs core proteins in the rescue of stressed hematopoiesis and restoration of blood homeostasis via m^6^A mRNA ribostasis. m^6^A modifications change with aging and remain largely unexplored in the contexts of the blood development (Gao et al., 2021). Our studies show the effect of SG-stored RNAs on tissue differentiation and how stress resolution can lead to RNA release, translation and facilitate regeneration, establishing a mechanism of SG-mediated cellular regeneration.

## Methods and materials

### Inducing Erythroid differentiation of hCD34+ cells

CD34+ Cells were differentiated using three-step protocol. The basal erythroid differentiation medium (EDM) contains Iscove’s Modified Dulbecco’s Medium (IMDM; ThermoFisher, Cat #12440061) and is supplemented with 5 mL of 100x stabilized L-glutamine (ThermoFisher, Cat #25030081), 330 μg/mL holo-human transferrin (Sigma, Cat #T4132), 10 μg/mL recombinant human insulin (Sigma, Cat #I9278), penicillin/streptomycin (Corning, Cat #30-002), 2 IU/mL heparin (StemCell, Cat #7980), and 5% Human serum (Sigma, Cat #H4522). In stage 1 (of 3) of the differentiation procedure (Days 0 to 7), cells were cultured in EDM1 media – EDM supplemented with hydrocortisone (StemCell, Cat #74142), Stem Cell Factor (SCF; PreproTech, Cat #300-07), Interleukin 3 (IL-3; PeproTech, Cat #200-03), and 3U/mL Erythropoietin (EPO; Epogen^®^ (epoetin alfa).. On day 4, four volumes of fresh EDM1 media were added to every 1 volume of cell culture. In stage 2 (of 3) of differentiation (Days 7 to 11), cells were resuspended and cultured in EDM2 media – EDM supplemented with SCF and EPO. Subsequently, in stage 3 (of 3) of the differentiation procedure (Days 11 to 18), cells were cultured in EDM3 media – EDM supplemented with solely EPO, and EPO was replenished every third-day post day 18.

### Hemin-induced erythroid differentiation of K562 cells

K562 cells were seeded at a density of 5 × 10^4^ cells/mL in vented cap polystyrene suspension flasks (Falcon), maintained at 37 °C, 5% CO_2_, and cultured in RPMI (Gibco) supplemented with 10% FBS and 1% 5,000 U/ml Pen-Strep with or without hemin (50 μM) for up to 4 days.

### Chromatin Immunoprecipitation (ChIP)

For Gata 1 and Gata 2 ChIP-seq experiments, we used Gata1 (Santa Cruz sc265X), Gata2 (Santa Cruz sc9008X) antibodies. ChIP experiments were performed as previously described with slight modifications (Choudhuri et al., 2020). Briefly, 20-30 million cells for each sample were crosslinked using 1/10 volume 11% fresh formaldehyde for 10 min at room temperature. The cross-linking was quenched by the addition of 1/20 volume 2.5M Glycine. Cells were lysed in 10 mL of Lysis buffer 1 (50 mM HEPES-KOH, pH 7.5, 140 mM NaCl, 1 mM EDTA, 10% glycerol, 0.5% NP-40, 0.25% Triton X-100, and protease inhibitors) for 10 min at 4C. After centrifugation, cells were resuspended in 10 mL of Lysis buffer 2 (10 mM Tris-HCl, pH 8.0, 200 mM NaCl, 1 mM EDTA, 0.5 mM EGTA, and protease inhibitors) for 10 min at room temperature. Cells were pelleted and resuspended in 3 mL of Sonication buffer for K562 and U937 and 1 mL for other cells used (10 mM Tris-HCl, pH 8.0, 100 mM NaCl, 1 mM EDTA, 0.5 mM EGTA, 0.1% Na-Deoxycholate, 0.05% N-lauroylsarcosine, and protease Inhibitors) and sonicated in a Bioruptor sonicator for 24-40 cycles of 30s followed by 1min resting intervals. Samples were centrifuged for 10 min at 18,000 g and 1% of TritonX was added to the supernatant. Prior to the immunoprecipitation, 50 mL of protein G beads (Invitrogen 100-04D) for each reaction were washed twice with PBS, 0.5% BSA twice. Finally, the beads were resuspended in 250 mL of PBS, 0.5% BSA and 5 mg of each antibody. Beads were rotated for at least 6 hr at 4^0^C and then washed twice with PBS, 0.5% BSA. Cell lysates were added to the beads and incubated at 4^0^C overnight. Beads were washed 1x with (20 mM Tris-HCl (pH 8), 150 mM NaCl, 2mM EDTA, 0.1% SDS, 1%Triton X-100), 1x with (20 mM Tris-HCl (pH 8), 500 mM NaCl, 2 mM EDTA, 0.1% SDS, 1%Triton X-100), 1x with (10 mM Tris-HCl (pH 8), 250 nM LiCl, 2 mM EDTA, 1% NP40) and 1x with TE and finally resuspended in 200 mL elution buffer (50 mM Tris-Hcl, pH 8.0, 10 mM EDTA and 0.5%–1% SDS) Fifty microliters of cell lysates prior to addition to the beads was kept as input. Crosslinking was reversed by incubating samples at 65^0^C for at least 6 hr. Afterwards the cells were treated with RNase and proteinase K and the DNA was extracted by Phenol/Chloroform extraction.

End repair of immunoprecipitated DNA was performed using the End-It End-Repair kit (Epicentre, ER81050) and incubating the samples at 25^0^C for 45 min. End-repaired DNA was purified using AMPure XP Beads (1.8X of the reaction volume) (Agencourt AMPure XP – PCR purification Beads, BeckmanCoulter, A63881) and separating beads using DynaMag-96 Side Skirted Magnet (Life Technologies, 12027). A-tail was added to the end-repaired DNA using NEB Klenow Fragment Enzyme (3’-5’ exo, M0212L), 1X NEB buffer 2 and 0.2 mM dATP (Invitrogen, 18252-015) and incubating the reaction mix at 37^0^C for 30 min. A-tailed DNA was cleaned up using AMPure beads (1.8X of reaction volume). Subsequently, cleaned up A-tailed DNA went through Adaptor ligation reaction using Quick Ligation Kit (NEB, M2200L) following manufacturer’s protocol. Adaptor-ligated DNA was first cleaned up using AMPure beads (1.8X of reaction volume), eluted in 100μl and then size-selected using AMPure beads (0.9X of the final supernatant volume, 90 μl). Adaptor-ligated DNA fragments of proper size were enriched with PCR reaction using Fusion High-Fidelity PCR Master Mix kit (NEB, M0531S) and specific index primers supplied in NEBNext Multiplex Oligo Kit for Illumina (Index Primer Set 1, NEB, E7335L). Conditions for PCR used are as follows: 98 °C, 30 sec; [98°C, 10 sec; 65 °C, 30 sec; 72 °C, 30 sec] X 15 to 18 cycles; 72°C, 5 min; hold at 4 °C. PCR enriched fragments were further size-selected by running the PCR reaction mix in 2% low-molecular weight agarose gel (Bio-Rad, 161-3107) and subsequently purifying them using QIAquick Gel Extraction Kit (28704). Libraries were eluted in 25μl elution buffer. After measuring concentration in Qubit, all the libraries went through quality control analysis using an Agilent Bioanalyzer. Samples with proper size (250-300 bp) were selected for next-generation sequencing using Illumina Hiseq 2000 or 2500 platform.

### hCD34+ and K562 cell CRISPR editing

For CRISPR editing RNPs containing 6μl of sgRNA (30 pmol/μl) and 1μl Cas9 (20 pmol/μl) were assembled in 13μl Nucleofector ^TM^ solution (LONZA) at room temperature for 30mins. For each experimental replicate, 1.5×10^5^ cells were transfected using Nucleofector ^TM^. CD34^+^ cells were then recovered in a medium containing STEM span + CC100 cytokine cocktail. Gene editing was confirmed using ~10,000 cells 24h post nucleofection by genomic DNA isolation, locus-specific PCR amplification, and sequencing. (See Tables 3 and 4 for details)

**Table 3.**
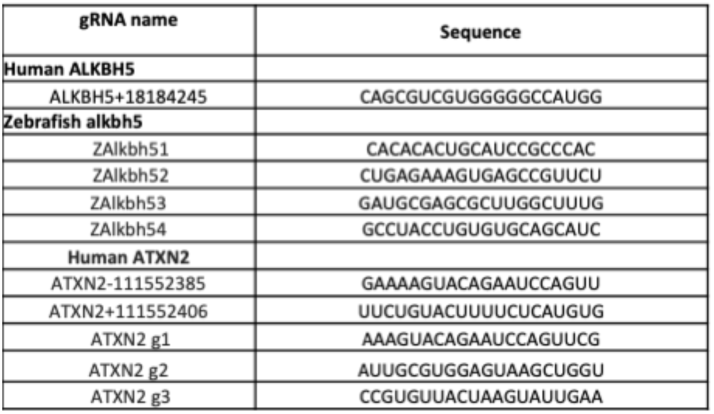
gRNA sequences used for editing *ALKBH5* and *ATXN2* genes.

**Table 4.**
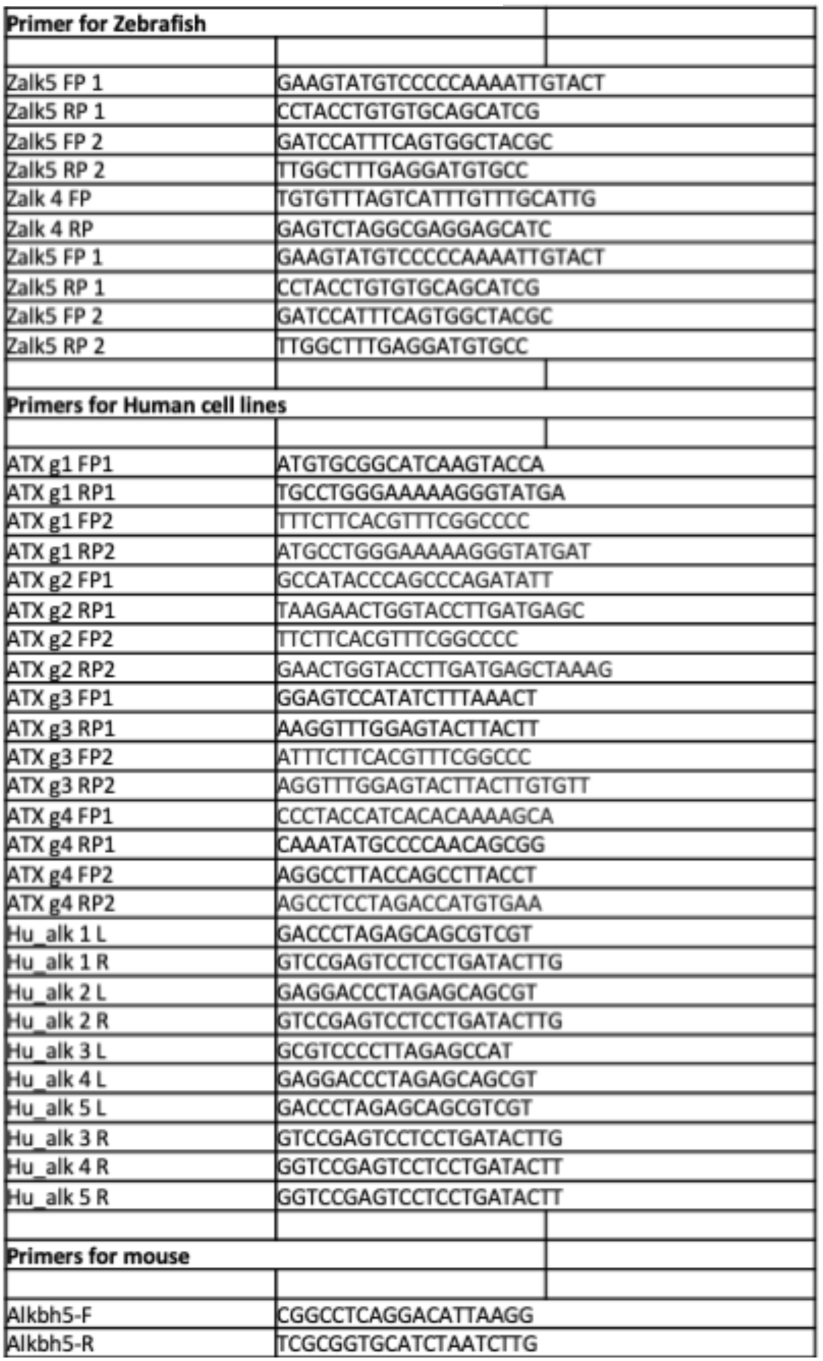
Primer sequences for Cas9-CRISPR editing experiments.

### Flow cytometric analysis of surface marker expression in hCD34+ and K562 cells

2.5 x 10^5^ cells per each condition were pelleted at room temperature and resuspended in 2ml FACS buffer (1X DPBS with 1% FBS). SYTOX^TM^ Blue dead cell stain (Invitrogen, Cat #S34857) was added at 1:2000 dilution and incubated in dark on ice for 5 minutes. Samples were then washed with 1 ml FACS buffer and pelleted at 4℃. Samples were then fixed in freshly prepared, chilled 4% paraformaldehyde in DPBS for 20 minutes at room temperature and subsequently washed with 1 ml DPBS and pelleted at 4℃. Cells were incubated with anti-CD235a (Glycophorin A, APC), and anti-CD71 (Transferrin Receptor 1, PE) for f30min at 4℃ at 60 rpm. At the end of incubation, samples were washed twice at 4℃ and resuspended in a final volume of 300 μl FACS buffer and analyzed on an MA900 Multi-Application cell sorter.

### Generation and genotyping of zebrafish mutant lines

The CRISPR-Cas9 system was used to generate mutations in *Alkbh5*.

gRNAs were designed using CHOP software and RNPs containing gRNA and Cas9 were injected into embryos at the 1-cell stage. For this 6μl sgRNA (30 pmol/μl) was incubated with Cas9 2NLS (20 pmol/μl; 3.22 mg/ml). The gene editing was validated using fin clips followed by genomic DNA isolation and locus-specific PCR amplification and sequencing.

To genotype mutants, DNA was extracted from clipped adult fin tissue through incubation; samples were incubated at 95°C in 50mM NaOH for 45 minutes, and subsequently vortexed and neutralized with 10% 1M Tris-HCl, pH 8. 10-50 ng genomic DNA was used for each PCR reaction.

### m^6^A sequencing and analysis

To understand mRNA targets, we probed for m6A marked transcripts during CD34^+^ and K562 cell erythropoiesis using sequencing (Dominissini et al., 2013; Lin et al., 2016). We followed a previously established protocol (Lin et al., 2016; Yang et al., 2022). The total RNA isolation (~300 micrograms) was fragmented to ~150 bp. A small part of this RNA ~10ng was kept aside as input and the rest was used for anti-m6A antibody-mediated immunoprecipitation to select m6A marked RNA fragments. cDNA is synthesized using reverse transcription of both input RNA and m^6^A immunoprecipitated RNA. Sequencing gives the list of m^6^A modified transcripts.

### m^6^A RIP-seq protocol

~50-60 million cells were used for each sample and total RNA was isolated using extraction Direct-zol RNA Miniprep Plus Kits (Cat #-R2072). Poly-A selection was performed using PolyATtract® mRNA Isolation Systems (Promega, Z5310). Poly-A selected RNA was resuspended in 15ul of nuclease free water. 1ul of ERCC RNA spike-in (0.1ng/ul) (Cat #4456740) was added and this was subjected to controlled fragmentation to get 150-200 bp long RNA fragments using a fragmentation buffer (See recipe section for details). 2 μl of 0.5 M EDTA was added to this mixture and RNA fragments were column purified (Zymo research Cat no-D4061) and eluted in RNAse free water. 4-5% of eluted RNA quantity of RNA was kept aside as input control for RNA sequencing. The remaining RNA was divided into two technical replicates for anti-m6A antibody-mediated pulldown to enrich for m6A-modified RNA fragments. m6A enriched RNA fragments and input RNA was processed following NEBNext® Ultra™ II RNA Library Prep Kit for Illumina protocol (NEB #E7775) and subjected to sequencing using Illumina.

The details of each step of the protocol are as follows:

**Fragmentation**- Assemble the following reaction using Poly-A selected RNA in an Eppendorf tube.

1. Assemble reaction with 15ul RNA, 2ul of 10X fragmentation buffer, 1ul Spike in RNA (3 diff. in vitro transcribed RNA) (0.1ng/ul), 2ul of water is assembled on ice.
2. Incubate for 5mins at 94℃ and immediately add 2 μl of 0.5 M EDTA to stop the reaction. Add 28ul of water and place it on ice for 5mins. Proceed to Column purification using Zymo research (D4061), elute sample with 210ul of water.
3. Store 10ul of RNA at −20℃ for input control. Take 100ul of RNA for each reaction and assemble m6A antibody-mediated immune precipitation (IP) reaction for two technical replicates. Incubate at 4℃ for 2hrs on a rotary shaker. Each reaction contains100ul of RNA sample, 5ul RNase inhibitor (40U/ul), 100ul 5X IP buffer, 5ul m6A antibody, 290ul of Water.
4. Prepare Pierce protein A/G Magnetic Beads (Thermo Scientific, #88803) by washing with 1x IP buffer for 2 times in a Magnetic stand. Remove all the buffer and dissolve beads in 1 ml of 1X IP buffer supplemented with BSA (0.5 mg/ml) and rotate it at 4℃ for 2hrs.
5. At the end of incubation wash beads on a magnetic stand with 1X IP buffer 2 times and add samples prepared in step 3. Incubate at 4℃ for 2hrs on a rotary shaker.
6. Place the tubes on a magnetic stand and perform four washes with 1X IP buffer. Remove all the buffer and add 100ul of elution buffer. Incubate at 4℃ for 1hrs on a rotary shaker.
7. Place the tubes on a magnetic stand and collect the supernatant in an Eppendorf placed on ice. Repeat step 6 and collect the supernatant. Finally, add 210ul of 1X IP buffer to magnetic beads, resuspend well and use the magnetic stand to collect the supernatant.
8. To 400ul supernatant, add 1ml EtOH and 40ul of 3M NaoAC to incubate at −80℃ for 12hrs. Spin at 13,000 rpm for 30min, at 4℃ to precipitate RNA. Discard the supernatant carefully. Dissolve RNA precipitate in 5ul of RNAse-free water.
9. Use both IP replicates and input control RNA from step 3 to prepare the library using NEBNext® Ultra™ II RNA Library Prep Kit. Use 15 PCR cycles for input control RNA and 19~20 cycles for IP duplicates at the end stage of library preparation.

**ZnCl2** (1 M): Dissolve 1.363 g of ZnCl2 in 10 ml of molecular biology–grade, RNase-free water. Store the solution at room temperature (22 °C) and use it within 18 months Composition

10X fragmentation buffer-

100ul of Tris-Cl (pH 7.0) (1M), 100ul of ZnCl_2_ (1M), and 800ul of Water

5X IP buffer

0.5mL of Tris-Cl (pH 7.4) (1M), 1.5mL of NaCl (5M), 0.5mL of NP-40 (10%), and 7.5mL Water

1X Elution buffer

90ul of 5X IP buffer, 150ul of m6A salt (20mM), 7ul RNase inhibitor and 203ul of Water

**m6A salt (20 mM)**: Dissolve 10 mg of m6A in 1.3 ml of molecular biology–grade, RNase-free water. Store aliquots of 150 μl at − 20 °C and use them within 12 months.

### m^6^A microarray for single nucleotide analysis (From Arraystar.com)

For the microarray, the total RNA was divided into two fractions. One of the fractions denoted as (Fraction1) “MazF-Digested” was treated with RNA endoribonuclease MazF. MazF cleaves the unmodified m6A sites. The Fraction 2, “MazF-Undigested” was mock-treated and serves as a control for modified and unmodified m6A sites. cRNAs (Complementary RNA) were synthesized with Cy5 label for Fraction 1 and Cy3 label for Fraction 2. Fraction 1 and 2 cRNAs were combined together and hybridized onto Arraystar Human m6A Single Nucleotide Array (8x15K, Arraystar). and were scanned in two-color channels by an Agilent Scanner G2505C. Images were analyzed using Agilent Feature Extraction software (version 11.0.1.1). Raw intensities of Fraction1 (Cy5-labelled) and Fraction2 (Cy3-labelled) were normalized with the average of log2-scaled Spike-in RNA intensities. “m6A site abundance” was calculated based on the probe signals having Present (P) or Marginal (M) QC flags in at least 3 out of 9 samples. “m6A site abundance” was calculated for the m6A site methylation amount based on the MazF-Digested (Cy5-labelled) normalized intensities. Differentially m6A-methylated sites between two comparison groups were identified by filtering with the fold change and statistical significance (p-value) thresholds. Hierarchical Clustering was performed to show the distinguishable m6A-methylation pattern among samples.

### Polysome analysis

#### Preparation of Samples for Polysome Profiling

CD34+ Cells were treated with 100 µg/ml cycloheximide (CHX) by directly adding it to the culture media for 15min prior to harvesting. Cells were pelleted at 600g for 5min and washed with PBS containing 100 µg/ml CHX. Cells were spun at 600 × g for 5 min at 4 °C or RT. Cell pellets were flash-frozen and stored at −80C to be used later for polysome profiling.

#### Density Gradient Ultracentrifugation and Polysome Profiling

Samples were thawed and lysed at 4C in polysome lysis buffer [10 mM Tris-HCl pH 7.4, 5 mM MgCl2, 100 mM KCl, 1% Triton X-100, 2 mM DTT, 100 μg/mL cycloheximide, cOmplete EDTA-free protease inhibitor cocktail (Roche), and 100 U/mL SUPERaseIn (Invitrogen)] trituration using a 26G needle, followed by incubation on ice for 10 min. Lysates were cleared by centrifugation at 10,000xg for 10 min at 4C. The resulting supernatant was transferred, mixed well, then layered onto a ~12 mL 10-50% sucrose gradient made in 10 mM Tris-HCl pH 7.4, 5 mM MgCl2, 100 mM KCl, 2 mM DTT, and 100 μg/mL cycloheximide. Samples were ultracentrifuged at 35,000 RPM using an SW-41 Ti rotor at 4C for 2.5 hr. Using a BioComp Piston Gradient Fractionator monitoring absorbance at 254 nm, gradients were analyzed and collected into ~24 fractions. For RNA sequencing, fractions were pooled into pre-polysome, monosome, and polysome fractions.

### Immunostaining and STED microscopy preparation of CD34+ and K562 cells

For each sample 50K cells were collected, rinsed with 500 ul PBS, and fixed by incubation using 100 ul of chilled 100% methanol for 5 minutes. Post-methanol fixation, cells were immediately rinsed with 500 ul PBS, and resuspended in 100 ul of 0.1% BSA in PBS. Cells were spun at 5 minute, 500 rpm, and medium acceleration (Thermo Shandon Cytospin 3 Centrifuge). Cells were incubated with primary antibodies (1:50 in 0.1% BSA), for 12 hours at 4°C on an orbital shaker. Cells were subsequently rinsed thrice with 200 ul PBS and then incubated in secondary antibodies (1:100 in 0.1% BSA) and a 20mM Hoechst 33342 Solution (1:100; Thermo Fisher, Cat #P36961) for 12 hours at 4°C on an orbital shaker. Cells were rinsed thrice with 200 ul PBS to remove unbound secondary antibodies. The stained cells were mounted using Diamond Antifade mounting medium (Invitrogen, Cat #P36961) and imaged using the Leica TCS SP8 Laser Scanning Confocal and Super Resolution microscope. (See Table 5 for details)

**Table 5.** Antibody details used for FACS analysis and microscopy experiments.

| Name | Company Details | Cat no |
| --- | --- | --- |
| ALKBH5 | Novus Biologics | NBP1-82188 |
| ATXN2 | sigma | HPA018295-25UL |
| GFP | abcam | ab13970 |
| PABP | abcam | ab21060 |
| N6-methyladenosine (m6A) antibody | abcam | ab151230 |
| CD235a | thermo | 17-9987-42 |
| PE Mouse Anti-Human CD71 | Biosciences | 555537 |
| G3BP | abcam | ab56574 |

### Phenylhydrazine induced anemia mouse model

The Alkbh5 (Embryonic knockout (KO))/B6 mouse model was applied and described in previous work (Shen et al., 2020). Briefly, Age-and-Sex matched Alkbh5 knockout mice or control mice were intraperitoneally injected with phenylhydrazine (114715, Sigma-Aldrich) at a dose of 100 mg/ kg body weight, and the erythroid level in peripheral blood was evaluated daily by CBC. For FACS analysis, the mice were injected with PHZ as described above. On day 2 post-injection, bone marrow cells and splenic cells were harvested and stained with Ter119 and CD71 antibodies for 30 minutes following by flow cytometry analysis in BD FACS CytoFLEX S.

### Phenyl hydrazine-induced acute anemia Zebrafish model

PHZ (Sigma-Aldrich) solution was used to induce hemolytic anemia. Adult zebrafish were treated as per the following protocol (McReynolds et al 2008). Adult fish (6–12 months, WT, het, and mutant) treated with PHZ and recovery were studied over a 8-day period. The fish were treated for 20 min with phenylhydrazine (0.003 mg/mL). Post PHZ exposure, the fish were rinsed with water for three minutes, four times. Whole kidney marrow was analyzed using flow cytometry.

---

**Supplementary Fig. 1:**
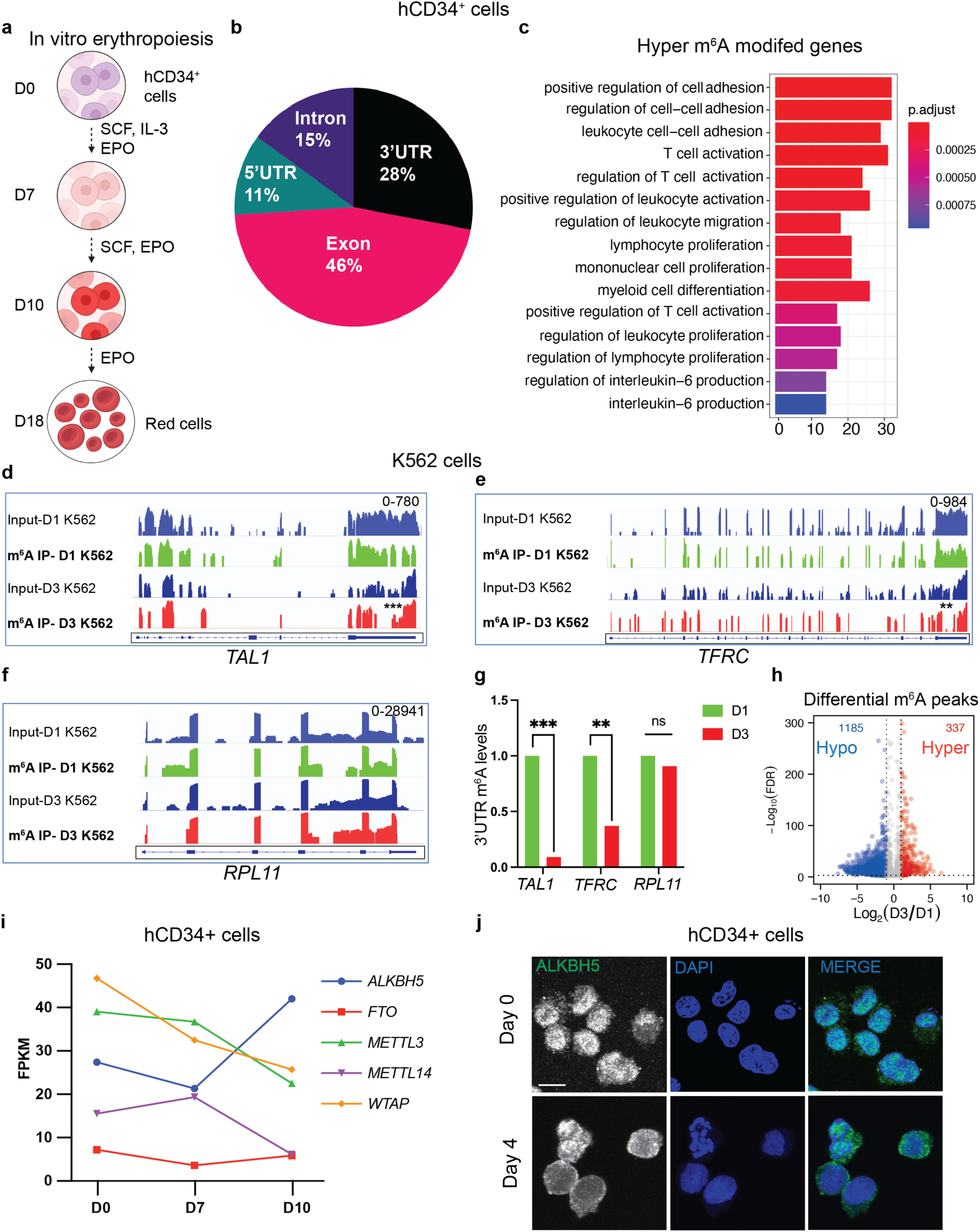
**a**, hCD34+ cells invitro erythropoiesis protocol. **b**, Pie chart graph showing the transcriptome-wide distribution of m^6^A marks during hCD34+ cell erythropoiesis. **c,** GO terms of genes with hyper m^6^A peaks. **d-h**, m^6^A analysis of K562 cell post hemin induction. K562 cells were subjected hemin induced erythropoiesis, and poly-A mRNA was subjected to m^6^A analysis using an anti-m^6^A antibody. Green and red plots represent m^6^A methylation marks along the gene for day 1 and day 3, respectively. Blue plots represent the total RNA input sequence. *TAL1*, *TFRC* and *RPL11* mRNA m^6^A levels are shown for day 1 and day3 post hemin induction. **g**, Graph showing quantification of m^6^A levels for *TAL1*, *TFRC* and *RPL11* genes. **h**, Differential m^6^A peaks during K562 cell erythropoiesis. Blue dots represent decreased m^6^A peaks (hypo-m^6^A), and red dots represent increased m^6^A peaks (hyper-m^6^A). Grey dots correspond to m^6^A peaks with no significant change. **i**, Transcript levels of *ALKBH5*, *FTO*, *METTL3*, and *METTL 14*, *WTAP* during hCD34+ cell erythropoiesis. **j**, hCD34+ cells immunostained using anti-Alkbh5 (green) and DAPI (blue) on Day 0 and Day 4 during erythropoiesis. p-values *** <0.001, ** <0.01 from unpaired t-test. H-hour, D-Day.

**Supplementary Fig. 2:**
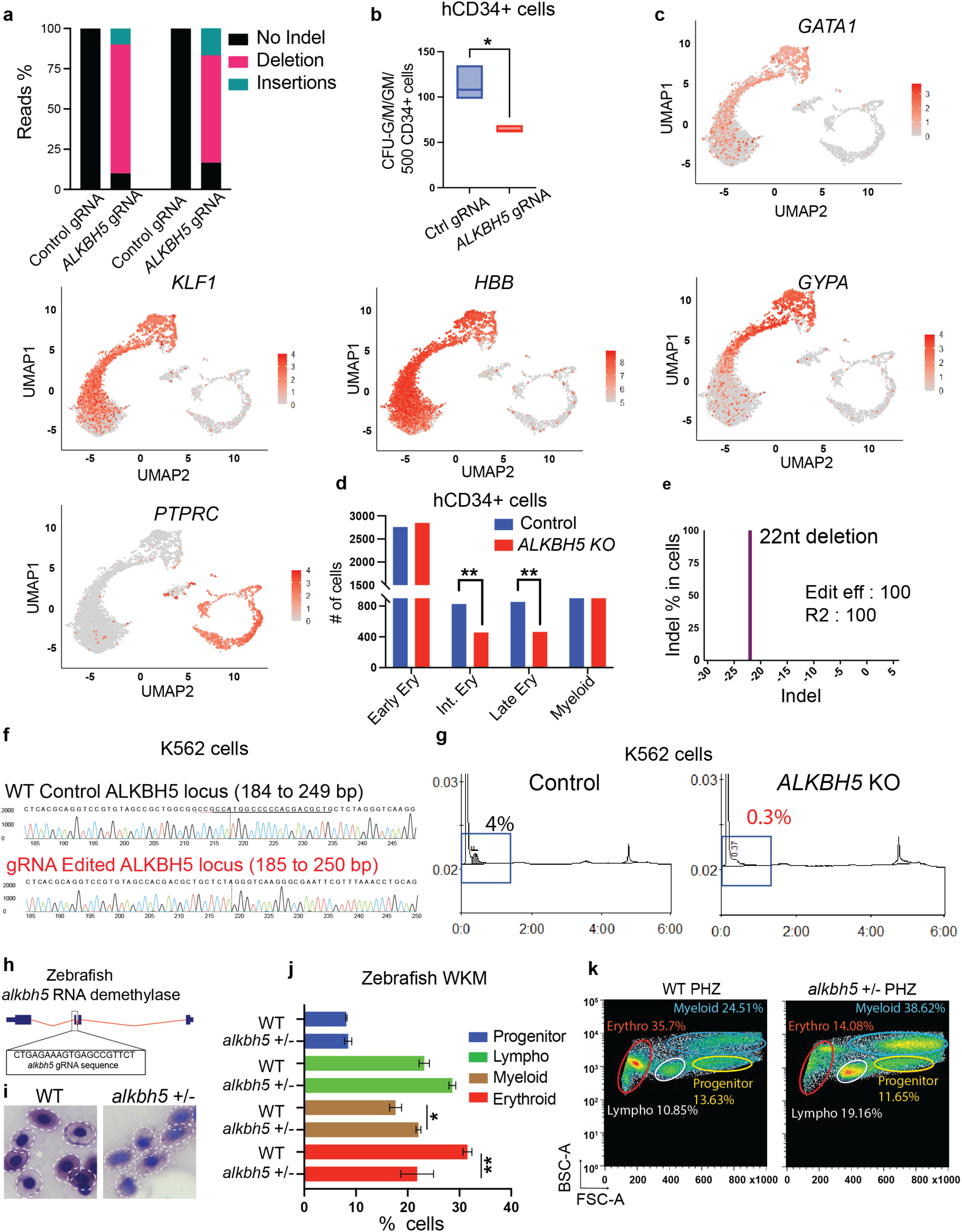
**a**, Graph showing indel frequency distribution for CD34+ cell CRISPR editing for Alkbh5. **b**, Quantification of CFU-G/M/GM per 500 CD34+ cells from methocult analysis. CFU-G/M/GM per 500 hCD34+ cells from methocult analysis. **c**, UMAP plots depicting gene clustering for erythroid cell and myeloid clusters. *GATA1, KLF1, HBB, GYPA*, marks erythroid cells and *PTPRC* shows myeloid cells from Methocult culture. **d**, Graph showing different cell types from UMAP plot. **e**, Graph depicting indel distribution corresponding to 22nt deletion in Alkbh5 locus. **f**, Sequencing tracks of human *ALKBH5* unedited and edited sequence from K562 cells using control gRNA and *ALKBH5* gRNA. **g**, HPLC analysis of K562 cells post hemin induction. K562 control and K562 *ALKBH5* KO cells were subjected to cell lysis at day 3 post hemin induction and HPLC analysis to measure gamma globin levels. **h**, Zebrafish *alkbh5* with gRNA targeting RNA demethylase sequence. **i**, Giemsa stain of peripheral blood showing red cells from WT and *alkbh5* +/−. **j**, Graph showing different blood cell type distribution in WT and *alkbh5* +/− fish. **k,** FACS plots of WKM blood cell types in WT and *alkbh5* +/− zebrafish on day 8 of recovery of phenylhydrazine-induced acute anemia. P-values ** <0.001 from unpaired t-test.

**Supplementary Fig. 3:**
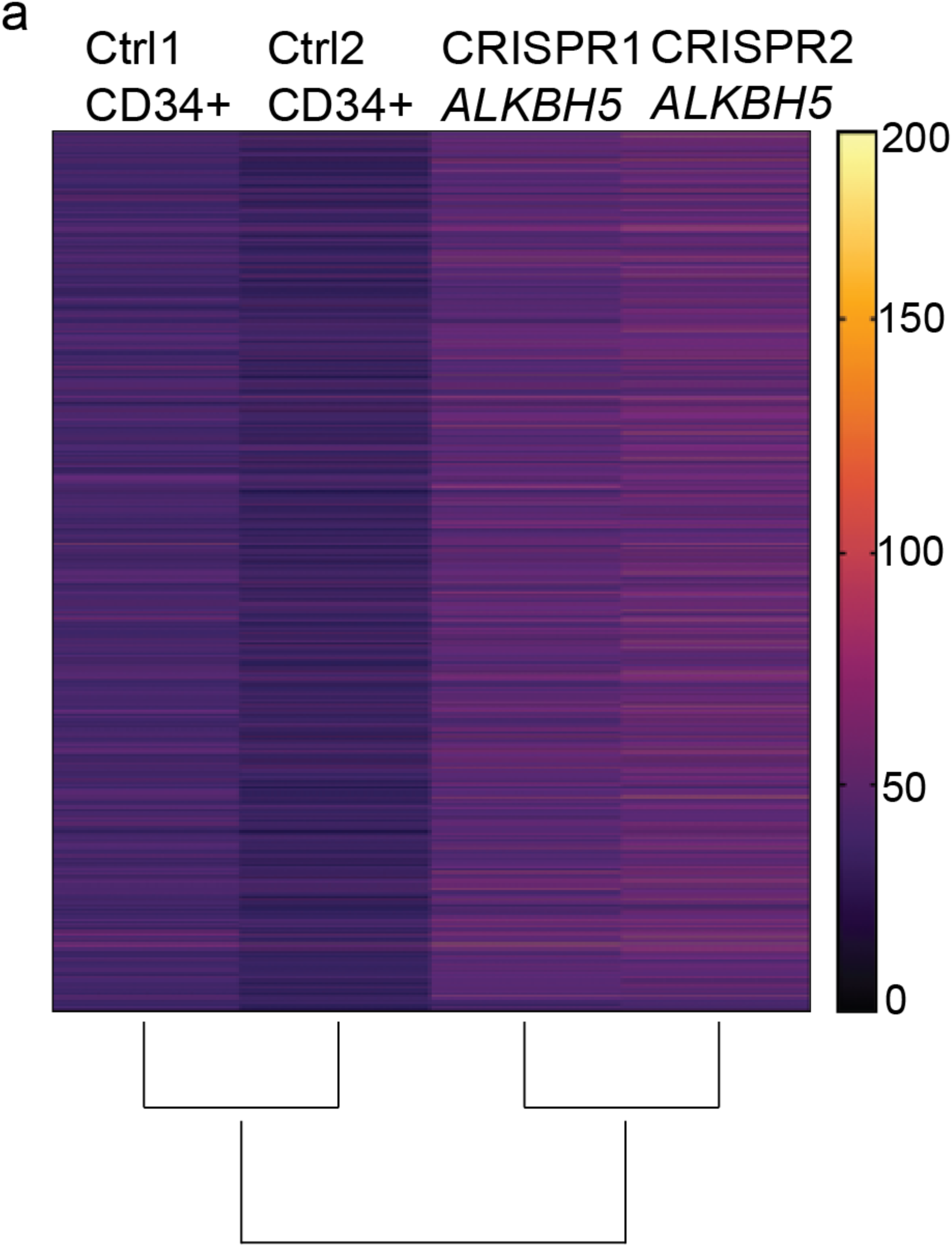
**a**, Hierarchical clustering heatmap of differentially m6A modified single m^6^A site. Transcriptome-wide m^6^A analysis of CD34+ cells edited for *ALKBH5* gene using CRISPR Cas9 technology. hCD34+ cells were nucleofected with control gRNA or *ALKBH5* gRNA+ Cas9, and m^6^A analysis was performed during erythropoiesis (day 7). The scale denotes methylation levels from 0-200 for differentially regulated 4800 genes.

**Supplementary Fig. 4:**
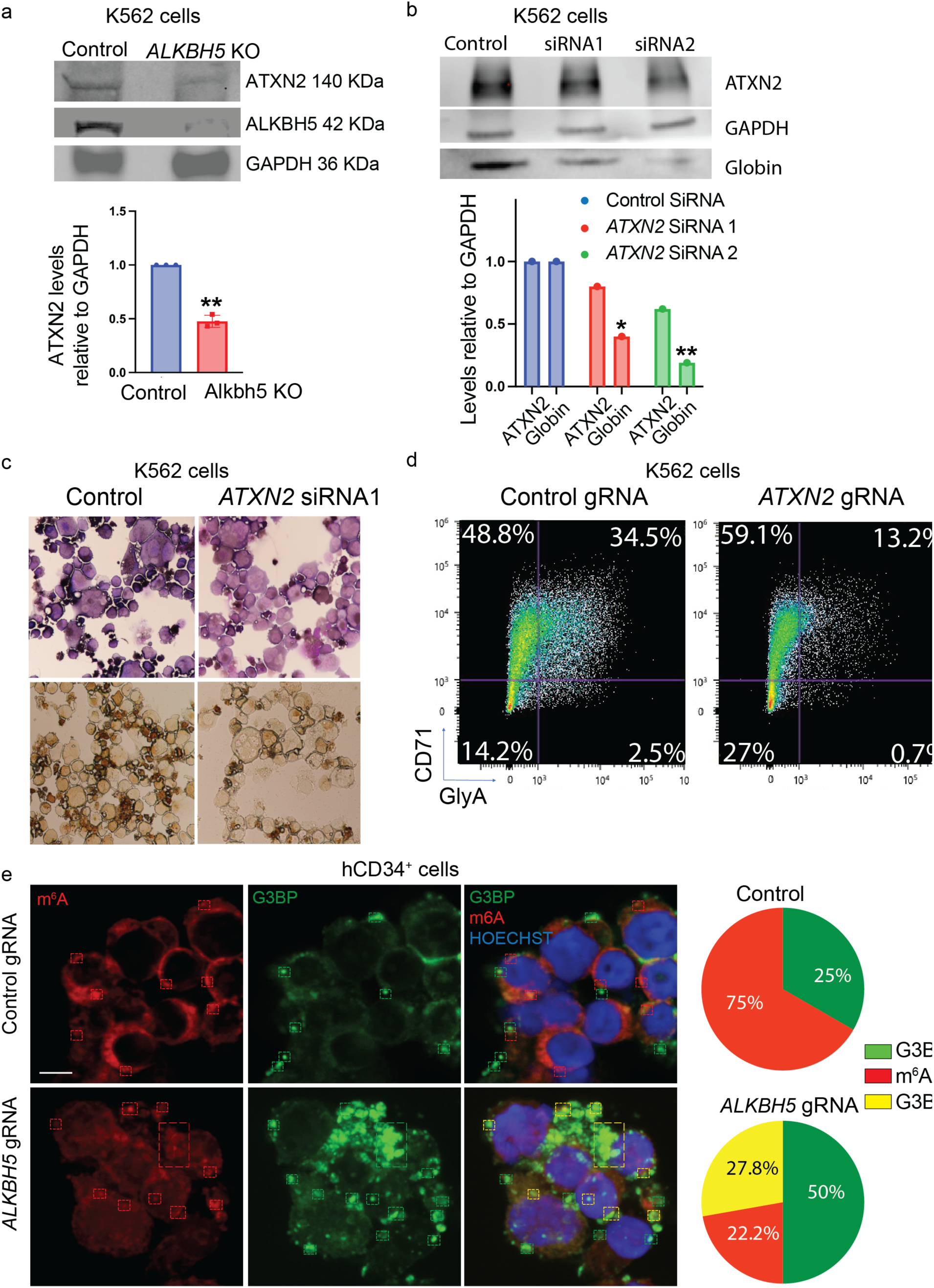
**a**, K562 *ALKBH5* KO cell analyzed for ATXN2 levels using western blot. K562 cells were probed on day 2 post hemin induction using anti-Alkbh5, anti-ATXN2 and anti-GAPDH antibodies. The graph depicts ATXN2 protein levels from western blot normalized to GAPDH protein levels. ** <0.01 from unpaired t-test. **b**, Western blot analysis of Control and *ATXN2* siRNA cells probed for gamma globin protein level. anti-ATXN2, anti-GAPDH, and anti-gamma globin antibodies were used to probe the levels of each protein. Quantification of the blot. GAPDH levels were used as loading control to normalize the quantification of ATXN2 and gamma globin levels. **c**, Benzedine and Giemsa stain analysis of K562 cells treated with control siRNA and *ATXN2* siRNA. K562 cells were subjected to siRNA transfection and hemin induction. Cells were imaged using benzidine and Giemsa stain on day 2 post hemin induction. n=3 **d**, FACS analysis of K562 control and *ATXN2* KO cells on day 2 post hemin induction, probed using Anti-TFRC, anti-GlyA antibodies. n=3 **e**, STED microscopy of hCD34+ cells during erythropoiesis. hCD34+ cells were subjected to control gRNA or *ALKBH5* gRNA. Cells were probed using anti-m^6^A, anti-G3BP antibodies, and Hoechst. Dotted rectangles show puncta labeled with anti-m^6^A and anti-m^6^A + anti-G3BP antibodies. Quantification of anti-G3BP, anti-m^6^A, G3BP+ m^6^A labeled puncta from n=3, N=20 cells, scale bar-5 μm, p-values *** <0.0001 from unpaired t-test.

**Supplementary Fig. 5:**
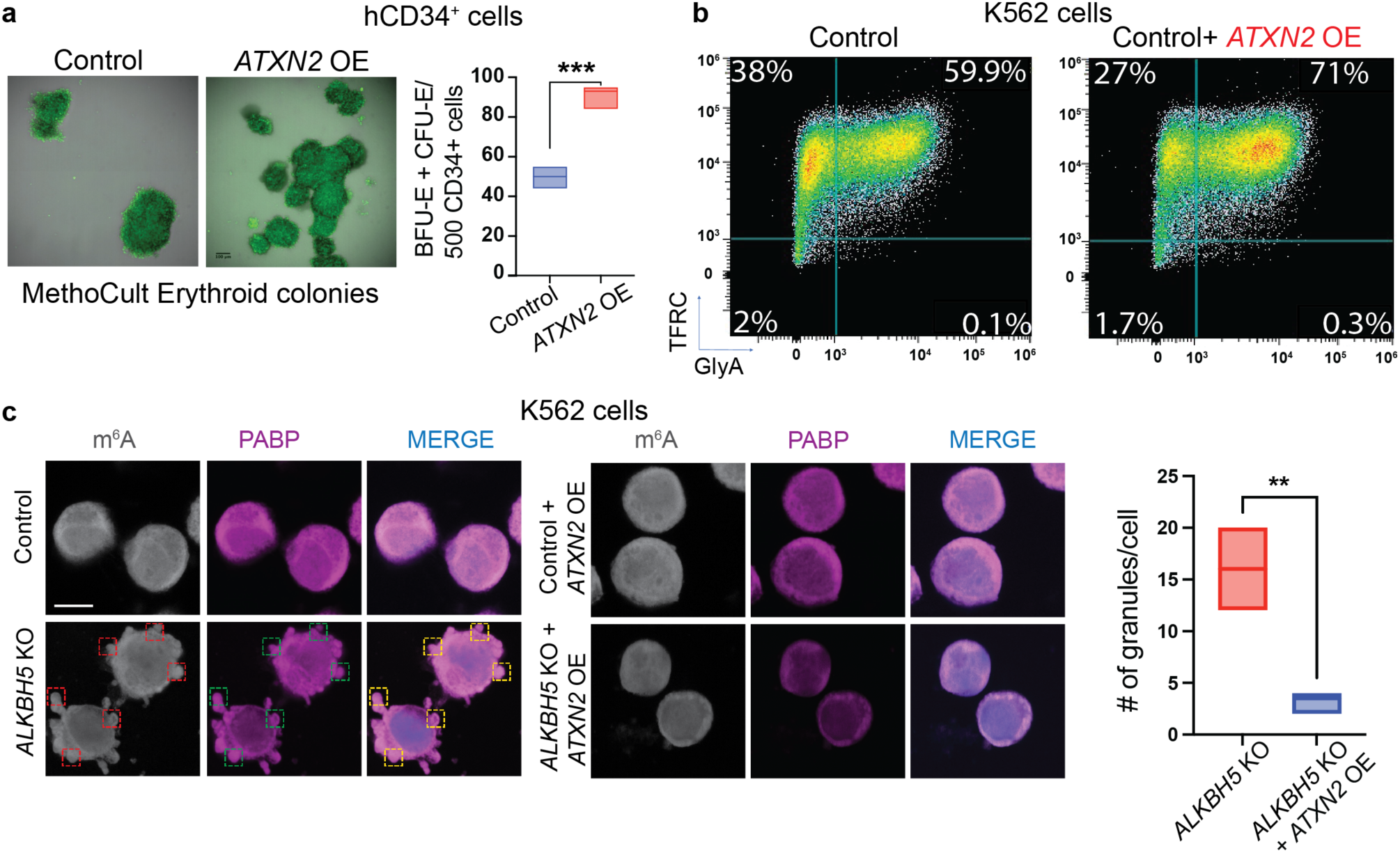
**a**, Methocult analysis of hCD34+ cells nucleofected with Control lenti vector or *ATXN2* overexpression (OE) lenti vector. Representative colony from each plate depicting erythroid colony size in control lenti vector or *ATXN2* overexpression (OE) lenti vector. n=3 hCD34+ donors, Scale bar-100 μm. Quantification of erythroid colonies per 500 hCD34+ cells from Methocult analysis. **b**, FACS analysis of day 2 hemin induced K562 cells transfected with control lenti vector or *ATXN2* overexpression (OE) lenti vector using anti-CD71 and anti-CD235a. **c**, K562 cells stained using an anti-PABP antibody. Control, *ALKBH5* KO, control + ATXN2 OE, and *ALKBH5* KO+ *ATXN2* OE K562 cells were subjected to hemin-induced erythropoiesis and processed on day 2 for STED microscopy. Dotted rectangles show puncta labeled with anti-m^6^A (red rectangles), anti-PABP (green rectangles), and anti-m^6^A + anti-PABP antibodies (yellow rectangles). Dotted rectangles show puncta labeled with anti-m^6^A (red rectangles), anti-PABP (green rectangles), and anti-m^6^A + anti-PABP antibodies (yellow rectangles). Quantification of PABP marked SGs in *ALKBH5* KO and *ALKBH5* KO+ *ATXN2* OE cells. Scale bar-10 μm, p-values *** <0.0001, ** <0.001 from unpaired t-test.

